# Malaria parasites undergo a rapid and extensive metamorphosis after invasion of the host erythrocyte

**DOI:** 10.1101/2024.09.03.610831

**Authors:** Aline Fréville, Flavia Moreira-Leite, Camille Roussel, Matthew R.G. Russell, Aurelie Fricot, Valentine Carret, Abdoulaye Sissoko, Matthew J. Hayes, Aissatou Bailo Dialo, Nicole Cristine Kerkhoven, Margarida Ressurreição, Safi Dokmak, Michael J. Blackman, Lucy M. Collinson, Pierre A. Buffet, Sue Vaughan, Papa Alioune Ndour, Christiaan van Ooij

**Author notes:** Centre for Ultrastructural Imaging King’s College London New Hunt’s House Guy’s Campus London SE1 1UL United Kingdom (Matthew R.G. Russell).

## Abstract

Within the human host, the symptoms of malaria are caused by the replication of malaria parasites within erythrocytes. Growth inside the erythrocyte exposes the parasites to the normal surveillance of erythrocytes by the host organism, in particular the clearance of erythrocytes in the spleen. Here we show that the malaria parasite *Plasmodium falciparum* undergoes a rapid, multi-step metamorphosis that transforms the invasive merozoite into an amoeboid-shaped cell within minutes after invading erythrocytes. This transformation involves an increase in the parasite surface area and is mediated by factors already present in the merozoite, including the parasite phospholipid transfer protein PV6. Parasites lacking PV6 do not assume an amoeboid form and instead are spherical and have a smaller surface area than amoeboid forms. Furthermore, erythrocytes infected with parasites lacking PV6 undergo a higher loss of surface area upon infection by *P. falciparum*, which affects the traversal of infected erythrocytes through the spleen. This is the first evidence that after invasion, the parasite undergoes a rapid, complex metamorphosis within the host erythrocyte that promotes survival in the host.

## INTRODUCTION

Malaria caused by the parasite *Plasmodium falciparum* remains a common cause of mortality and morbidity^1^. All symptoms of the disease are caused by infection of host erythrocytes, in which the parasite undergoes a well-defined 48-hour developmental process. The first half of the intraerythrocytic cycle is commonly referred to as the ‘ring’ stage, as the parasites assume a ring-shaped morphology on Giemsa-stained smears. The parasite then matures to what is referred to as the ‘trophozoite’ form; at this stage the infected erythrocyte is less deformable and adheres to the endothelial cells lining the capillaries, thereby sequestering it from the circulation. The parasite subsequently initiates DNA replication and undergoes nuclear division during the final ‘schizont’ stage, culminating in the formation and release of the infectious progeny, the ‘merozoites’. After their release from the host erythrocyte, in a process called egress^2,3^, merozoites bind to and invade erythrocytes to propagate the infection.

As erythrocytes infected with ring-stage parasites are present in the peripheral circulation, the parasites need to adapt to an important host defence against blood-borne pathogens: removal of erythrocytes in the spleen. In the red pulp of this organ, blood is filtered through extremely narrow passages (∼0.65 µm) called interendothelial slits (IES). Passage through the IES requires that erythrocytes are highly deformable; erythrocytes with decreased deformability, such as senescent and phenotypically altered erythrocytes, are removed^4–6^. The importance of the spleen for the control of malaria infection is evident in asplenic individuals; whereas parasites present in the circulation of individuals with a spleen are mostly ring-stage parasites, trophozoites and schizonts are readily observed in the circulation of splenectomized individuals^7^. However, even erythrocytes containing ring-stage parasites are susceptible to removal by the spleen^8^, with the most ‘spherocytic’ (spherical) erythrocytes preferentially filtered out^9^.

Although the development of malaria parasites inside the host erythrocyte has been studied for well over a century, the events leading to the establishment of the parasite inside the host cell have not been investigated in detail, owing to the difficulty in synchronizing the invasion events and, until recently, in genetic manipulation of the parasites. Hence, how the parasite transforms from the extracellular form to the intracellular ‘ring’ form is not well understood. Even though this stage is referred to as a ‘ring’, it is widely appreciated that *P. falciparum* parasites can assume an amoeboid (or dendritic) shape during the first part of the intraerythrocytic cycle^10–15^. However, only two studies have investigated this form in some detail, which revealed that parasites can assume an amoeboid form early during the intraerythrocytic cycle and freely interconvert between amoeboid and spherical forms^12,15^. The amoeboid forms can have multiple, motile ‘limbs’^11–14^, but neither the mechanism and timing of formation of the amoeboid shape after invasion nor its role in the survival of the parasite in the host have been established. Inhibiting amoeboid formation with jasplakinolide, which affects actin dynamics, did not affect the replication of the parasite^12^, indicating that *in vitro*, transition to this form is not required. However, the effect of jasplakinolide does indicate that the transition to the amoeboid form is an active process. In this study, we investigated the mechanism of the parasite transition to the amoeboid form after invasion, its role in parasite growth and the involvement of PV6, a parasite phospholipid transport protein, a mutant of which has previously been shown to form aberrant rings in Giemsa-stained smears ^16–18(ref)^. Combining volume (3D) electron microscopy (serial block-face scanning electron microscopy (SBF-SEM)), tight synchronization of parasite erythrocytes invasion and conditional mutagenesis, we observed that upon invasion, rather than passively transitioning to a ‘ring’ shape, *P. falciparum* parasites undergo a rapid and complex metamorphosis from the merozoite to the amoeboid form.

This process involves a parasite-driven expansion of its surface area that occurs in at least two distinct steps. Importantly, the parasite phospholipid transfer protein PV6 (PFA0210c/PF3D7_0104200, also referred to as PfSTART1^18,19^), is essential to progress to the amoeboid form, affecting the second growth step. PV6-mediated modifications of the erythrocyte are important for the survival of the parasite in the host, as mutants lacking this protein are less able to traverse the spleen. Our work describes in detail and provides mechanistic insight into the transformation of malaria parasites from merozoites to the amoeboid shape and reveals that the changes in the erythrocyte are important for survival in the host.

## RESULTS

### Merozoites rapidly transform to the amoeboid form after invasion

To establish when after invasion parasites transform into the amoeboid form, we observed parasites fixed at specific times after invasion. In these samples, parasites were detected in a variety of shapes [Fig 1A]; as parasites interconvert between amoeboid and round forms, these different shapes likely represent intermediates of the interconversion process. Note that in this study, all non-round parasites, including those with a single limb (flask-shaped) and square parasites, were scored as amoeboid [Fig 1A]. To ensure that invasion was highly synchronous, we used reversible egress inhibitors (Compound 2 and ML10) that allow egress (and hence invasion) of the parasites to be precisely timed by removal of the inhibitor^20–23^; arrested parasites egress and invade approximately 15 minutes after removal of the inhibitor^22^ [Supp Fig 1A]^20–23^. Amoeboid forms could be detected as early as 20 minutes after removal of the egress inhibitor, suggesting that parasites can assume an amoeboid form within five minutes after invasion [Fig 1B]. Live-cell imaging confirmed this observation; when parasite invasion was viewed live, the average time from invasion to amoeboid formation was approximately 17.5 minutes [Figs 1C and 1D; see Supp Fig 1B for an additional example of an invading parasite], with several parasites transitioning to the amoeboid form within ten minutes. The slight difference in the apparent rate of conversion to the amoeboid form in the two experiments may reflect that the observation of the live parasites may have negatively affected the transformation. Hence, the conversion from merozoite to the amoeboid form after invasion is very rapid, most often occurring within the first 20 minutes after invasion.

**Figure 1.**
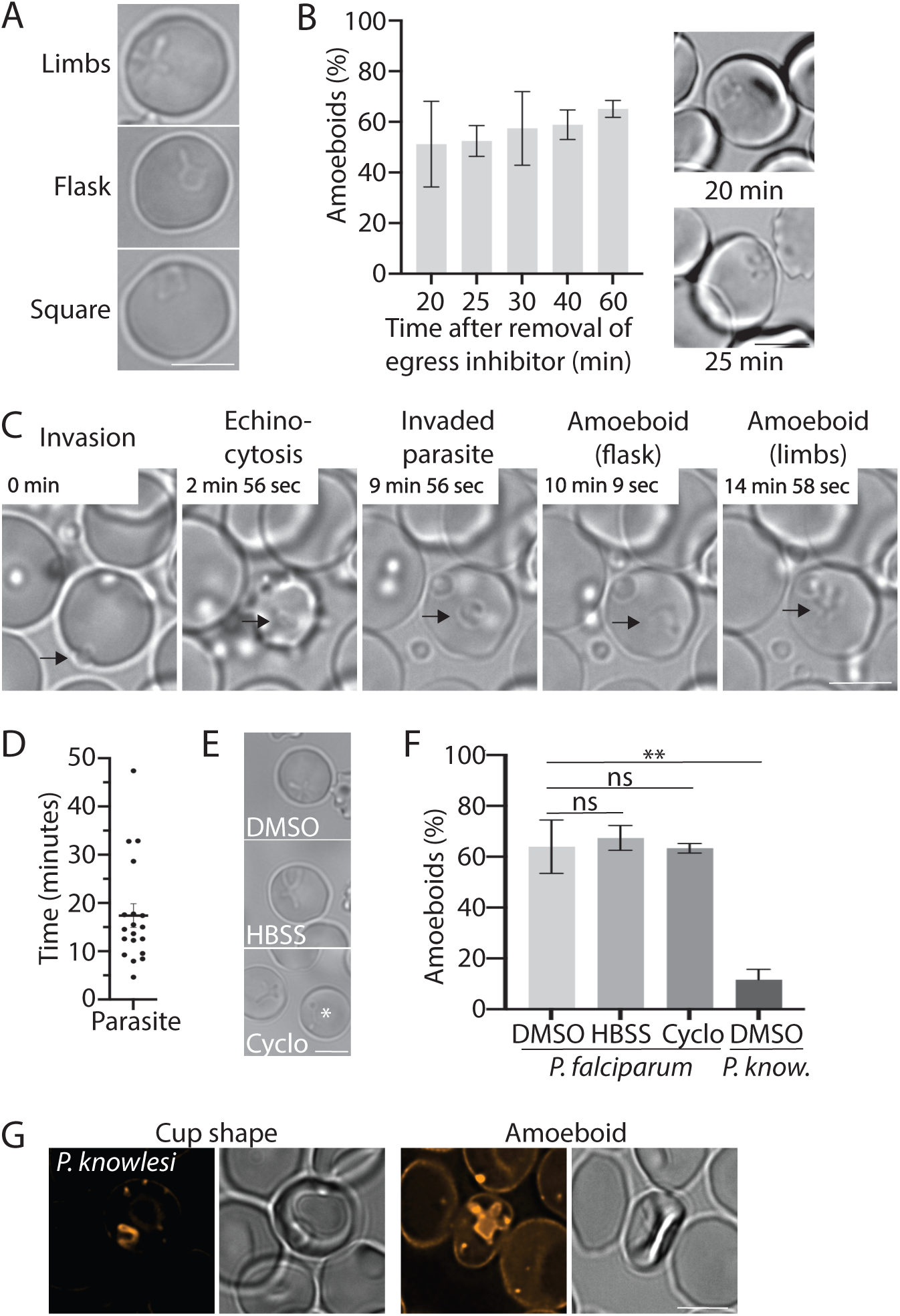
Amoeboid formation of *Plasmodium* parasites during the early erythrocytic cycle. **A.** Morphologies of *P. falciparum* parasites that were scored as amoeboid forms. **B.** Timing of amoeboid formation after invasion. Parasites were fixed at the indicated time after removal of ML10 and amoeboid formation was subsequently scored by microscopy. Results shown are the combination of two biological replicates. Errors bars indicate SD. Panels on the right are examples of amoeboid forms detected at the indicated time. **C.** Live-cell imaging of amoeboid formation. The frame rate was one frame/second, time starting from invasion is indicated in each panel. The arrows indicate the parasite. **D.** Timing of amoeboid formation in 19 parasites observed from the moment of invasion by video microscopy. Errors bars indicate SEM. **E and F.** Amoeboid formation of *Plasmodium falciparum* parasites after invasion in the presence of either DMSO in cRPMI (DMSO), Hanks’ Balanced Salt Solution (HBSS) or 5 µM cycloheximide (Cyclo) in cRPMI 2 hours after removal of ML10 and *Plasmodium knowlesi* 1-3 hours after removal of ML10. (E) Representative images of live *Plasmodium falciparum* parasites. The (*) indicates an oddly shaped elongated parasite that was occasionally detected in the cycloheximide-treated samples. (F) Amoeboid fornation in the indicated condition. Data represent three biological replicates of a minimum of 100 parasites (*P. falciparum)* and two biological replicates of *Plasmodium knowlesi* of at least 50 parasites (two-tailed t-test: ns: not significant; **P < 0.01, errors bar indicates SD)**. G.** Live-cell fluorescence imaging of *Plasmodium knowlesi*-infected erythrocytes stained with C5-Bodipy-ceramide 2 hours after removal of egress inhibitor. All scale bars represent 5 µm.

### Merozoites contain all the parasite components for the transformation to the amoeboid form

To gain insight into the mechanism underlying transformation to the amoeboid form, wildtype parasites were allowed to invade erythrocytes in the presence of either Hanks’ Balanced Salt Solution (HBSS), which contains only salts and glucose, or culture medium supplemented with the protein synthesis inhibitor cycloheximide. When evaluated by live microscopy two hours after removal of egress inhibitor, neither treatment had significantly affected the transformation to amoeboids, with ∼65% of intraerythrocytic parasites assuming an amoeboid form under all conditions [Figs 1E and 1F], indicating that neither external nutrients nor *de novo* protein synthesis are required for the transformation of parasites into amoeboid forms following invasion. Hence, merozoites appear to contain all the necessary parasite components to drive this transformation.

To determine whether other *Plasmodium* species form amoeboids, we observed *Plasmodium knowlesi* parasites 1-3 h after invasion. Amoeboid forms were detected, albeit at a much lower frequency compared to *P. falciparum* (∼10% vs ∼65%) [Fig 1F]. Instead, most *P. knowlesi* parasites formed a cup shape [Fig 1G], as has been reported previously^24^. The *P. knowlesi* amoeboid parasites appeared to be as flexible and able to interconvert between amoeboid and spherical forms as *P. falciparum* parasites^12^ [Supp Movies 1 and 2].

### Parasites lacking PV6 cannot assume an amoeboid form

We and others have previously shown that parasites lacking PV6 or parasites in which PV6 is inhibited remain small after invasion compared to wildtype and untreated parasites^16–18^. However, previous studies did not examine whether this reflects a collapse of the parasite or whether the parasite lacking PV6 activity simply fails to grow. To address this, we used a rapamycin-inducible *pv6* mutant in which *pv6* is excised after addition of rapamycin, allowing the phenotype of parasites lacking PV6 to be analyzed, even though PV6 is essential for parasite proliferation^16^. When trophozoites were treated with rapamycin approximately 30 hours after invasion, they matured seemingly normal prior to egress – they produced the same number of merozoites, had the same DNA content at the schizont stage and survived similarly to DMSO-treated parasites and the resulting merozoites invaded at the same rate [Supp Figs 2A-2D]. In contrast, the development of the parasites after invasion was severely affected. Measurement of the area of recently invaded parasites in Giemsa-stained smears revealed no difference in the size of wildtype (DMSO-treated) and mutant (rapamycin-treated) parasites 20 minutes after removal of the egress inhibitor, at which point the parasites had invaded the host cell approximately 5-10 minutes prior^22^. Thirty minutes after removal of the egress inhibitor, mutant parasites were significantly smaller than the wildtype and did not form standard ring forms on Giemsa-stained smears [Figs 2A and 2B; Supp Fig 2E, Supp Table 1]. The size of the parasites lacking PV6 did not increase further, indicating that these parasites do not develop any further after establishing themselves in the host erythrocyte. Hence, PV6 most likely functions immediately after invasion. Despite their small size, parasites lacking PV6 remained viable for an extended period after invasion, as indicated by positive MitoTracker staining [Supp Fig 2D]; similar survival of parasites was observed after treatment with a PV6 (PfSTART1) inhibitor^18^. This makes PV6 one of the first proteins known to be required immediately following invasion of the host cell, along with the inner membrane complex protein IMC1g and an unidentified Plasmepsin V target protein, both of which also affect parasite development at this stage^17,25^.

**Figure 2.**
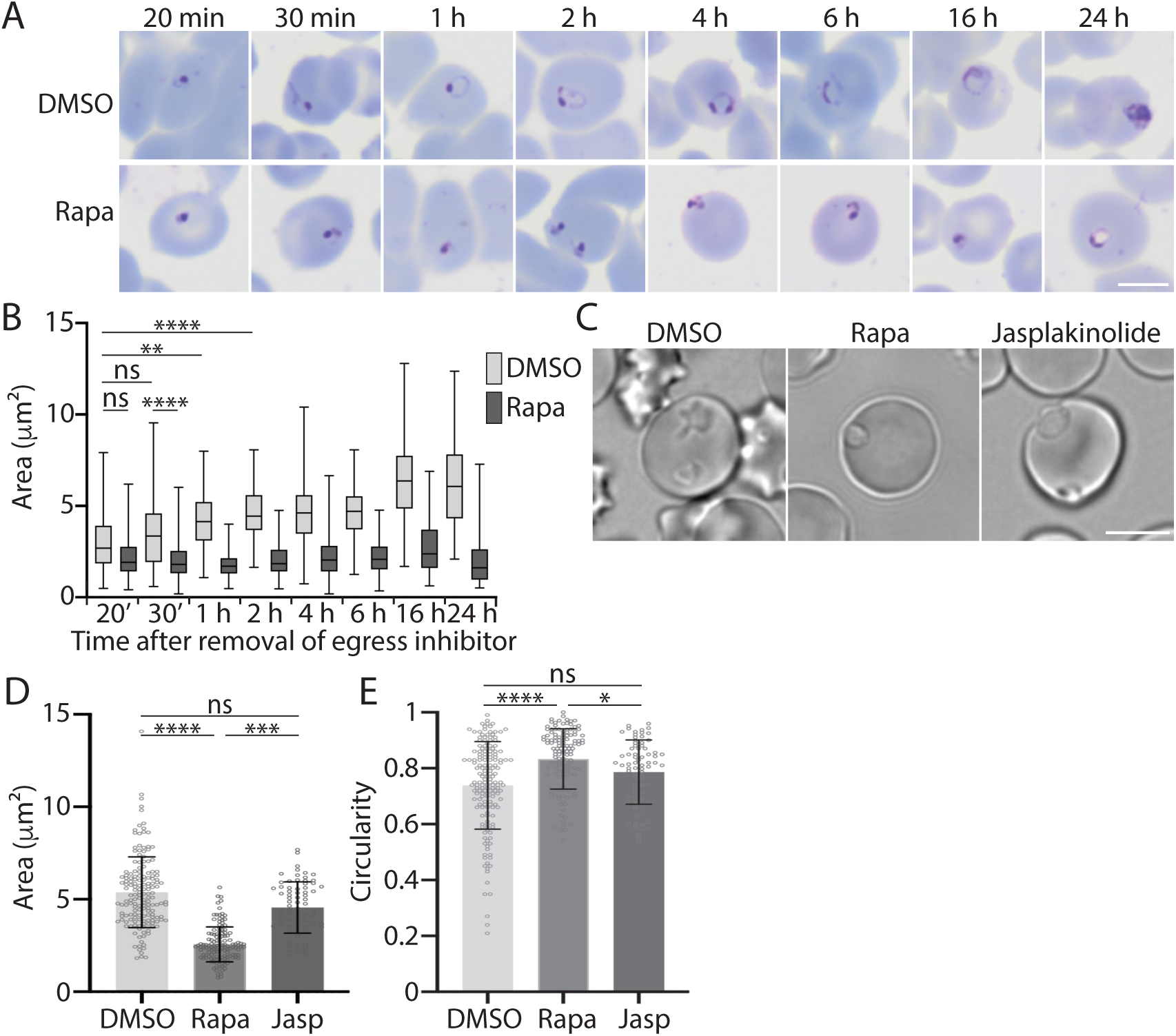
**Development of wildtype parasites and parasites lacking PV6 at early time points post-invasion**. **A.** Giemsa-stained smears of PV6 DiCre (PfBLD529) parasites treated with DMSO or rapamycin (Rapa). Development of synchronized parasites was examined in the cycle following DMSO or rapamycin treatment at the indicated time after removal of egress inhibitor (see Supp Fig 1 for outline of experiment). Scale bar represents 5 µm. **B.** Size (area) of DMSO and rapamycin-treated PV6-diCre parasites. Data represent three biological replicates with a minimum of 20 parasites per sample. Whiskers indicate range from minimum to maximum. Significance was determined using Kruskal-Wallis test: ns: not significant, *P < 0.05, ***P < 0.0001. For additional statistical analysis, please see Supp Table 1. **C**. Live-cell imaging of PV6-DiCre (PfBLD529) parasites treated with DMSO or rapamycin in the previous cycle or with jasplakinolide for one hour prior to imaging. Parasites were imaged 2 hours after removal of ML10. Scale bar represents 5 µm. **D.** Quantification of the size (area) of the parasites in the experiment presented in panel C. Data are based on measurement of at least 20 rings in each of the three biological replicates performed. Rapa: rapamycin-treated; Jasp: jasplakinolide-treated (Kruskal-Wallis test: ns: not significant, ***P < 0.001, ****P < 0.0001 ns: not significant; **P < 0.01, Errors bars represent SD). **E.** Analysis of the circularity of the parasites in the experiment presented in Panel C. Rapa: rapamycin-treated; Jasp: jasplakinolide-treated (Kruskal-Wallis test: ns: not significant, *P < 0.05, ****P < 0.0001, Errors bars +/-SD)..

Importantly, observation of wildtype and mutant parasites by live microscopy showed that wildtype parasites assumed an amoeboid form after invasion but that parasites lacking PV6 did not – these parasites were smaller and more spherical [Figs 2C-2E]. As treatment of wildtype parasites with jasplakinolide (which prevents amoeboid formation^12^), affected the shape but not the area, of wildtype parasites [Figs 2C-2E], we conclude that the smaller size of the mutant parasites does not reflect an inability to develop into the amoeboid form but rather an inability to increase in size. Hence, PV6 is required to progress to the amoeboid form and promotes the size increase of the parasite.

### The transition from merozoite to the amoeboid form is a two-step process promoted by a rapid increase in surface area

To investigate the changes in parasite shape in detail, we applied SBF-SEM – a 3D volume electron microscopy technique – to obtain series of hundreds of consecutive EMs sections separated by 50 or 70 nm (Supp movies 3-5). These data were then used to generate 3D models of parasites and erythrocytes. We focused on two time points: 20 minutes and two hours after the removal of egress inhibitor and compared wildtype parasites with parasites lacking PV6 at these times. Flattened wildtype parasites resembling the amoeboid form could already be detected 20 minutes after removal of the egress inhibitor, although only few parasites had formed multiple limbs at this time-point [Fig 3A, see Supp Fig 3 for additional views and Supp Fig 4A for additional models, Supp movie 3]. Two hours after removal of the egress inhibitor wildtype parasites had assumed various shapes, some with multiple limbs, consistent with the finding that these parasites are very motile and actively interconvert between amoeboid and spherical forms^12^. Interestingly, all limbs of the amoeboid parasites appeared to be attached to a central section of the parasite; no further bifurcation of the limbs was detected in any parasite, which potentially indicates that the parasite contains one central organizing center from which the limbs extend.

**Figure 3.**
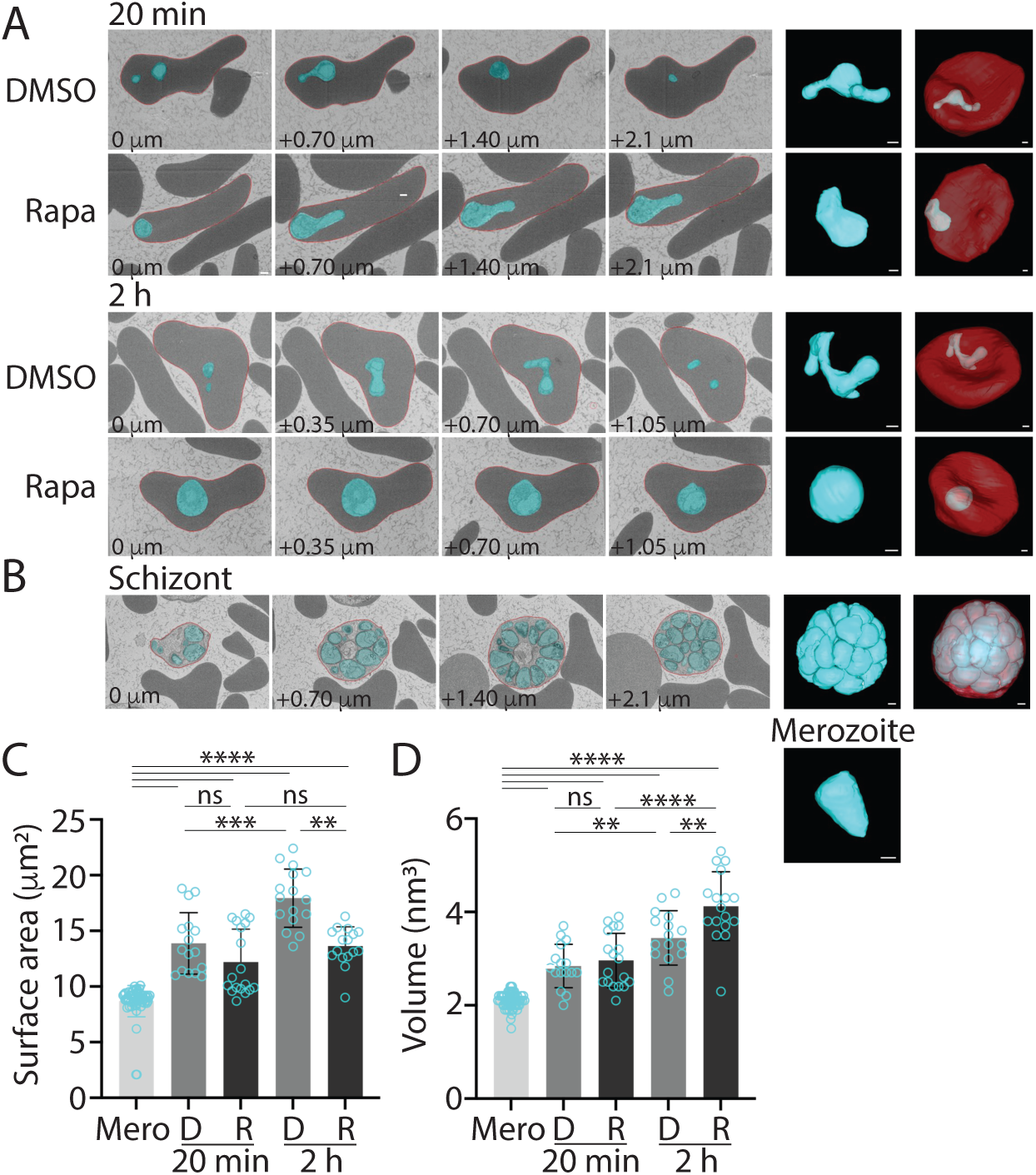
Amoeboid formation is accompanied by a large increase in surface area and requires PV6. **A.** SBF-SEM slices (sections) and surface renderings of PV6-DiCre (PfBLD529) parasites treated with DMSO and rapamycin 20 minutes and 2 hours after removal of egress inhibitor, illustrating the shape and positioning of the parasite (cyan) within the erythrocyte (red). Indicated in the panels is the distance between the sections. The panel second from right shows a 3D model of the parasite shown in the sections on the left-hand side. On the far right is shown the parasite modeled inside the erythrocyte. Scale bar represents 500 nm. **B.** Modeling of schizont and individual merozoite. **C and D.** Surface area (C) and volume (D) of merozoites (Mero) and DMSO-treated (D) and rapamycin-treated (R) parasites at the indicated time after removal of egress inhibitor. Data represents the measurement of at least 15 parasites. The Mann–Whitney U test was performed for statistical analysis: ns-not significant; *P <0.05; **P <0.01; ***P<0.001; ****P<0.0001. See Supp Fig 3 for additional views of these parasites and Supp Fig 4 for additional models.

**Figure 4.**
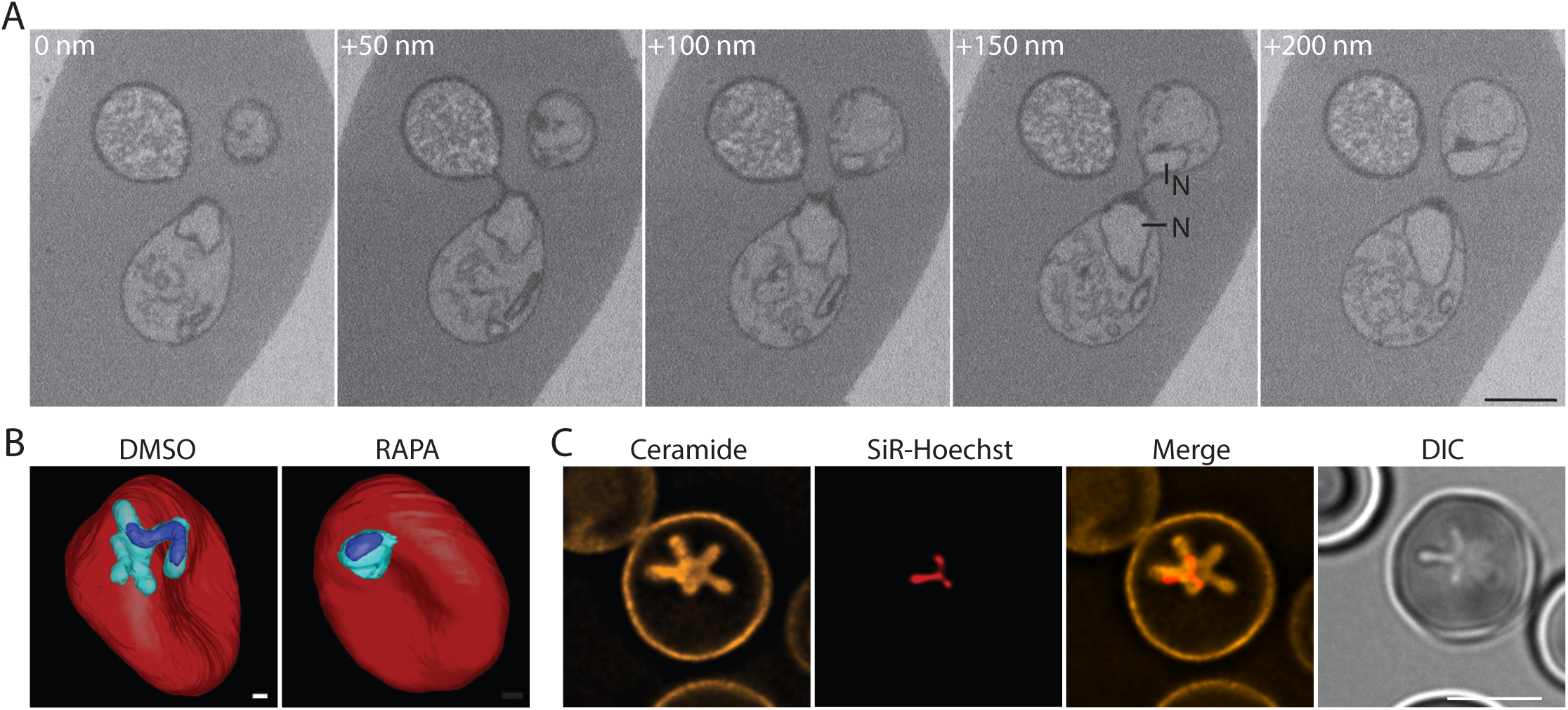
Connections between limbs of the amoeboid form are very narrow and can separate the nucleus into lobes. **A.** Consecutive SBF-SEM sections, showing three limbs of a parasite and the thin connections between them. Numbers in upper left-hand corner of the panels indicate distance between the sections, N indicates the nucleus and the scale bar represents 500 nm. **B.** Three-dimensional models of infected erythrocytes (red) two hours after removal of egress inhibitor, showing the nucleus (dark blue) in a wildtype (DMSO) parasites and a parasite lacking PV6 (RAPA); parasites are shown in cyan. Scale bar represents 500 nm. **C.** Live-cell imaging of an infected erythrocyte stained with ceramide, the DNA dye SiR-Hoechst, the merge of the ceramide and SiR-Hoechst images and imaged using DIC (right) two hours after the removal of egress inhibitor. The imperfect alignment of the ceramide and SiR-Hoechst staining is the result of the movement of the parasite during imaging. Scale bar represents 5 µm.

In contrast to wildtype parasites, parasites lacking PV6 did not possess limbs or assume an amoeboid shape. These parasites were more spherical than wildtype parasites but did assume somewhat extended forms 20 minutes after removal of egress inhibitor. However, two hours after egress inhibitor removal, these parasites had become nearly completely spherical [Fig 3A; Supp Fig 3 for additional views and Supp Fig 4A for additional models, Supp movie 5], indicating that PV6 is required for the progression to the amoeboid form.

The presence of late-stage schizonts containing fully formed, segregated merozoites in the samples obtained two hours after removal of the egress inhibitor allowed individual merozoites to be modelled as well [Fig 3B]. Furthermore, the infected erythrocyte could be modelled, revealing that the parasite occupies most of the erythrocyte [Fig 3B, Supp Fig 3].

The 3D models of the invaded parasites and the merozoites allow the surface area of the parasites to be determined and the growth of the parasite to be quantitated. This revealed that the surface area of wildtype parasites increased between the 20-minute and the two-hour time points, consistent with the analysis of Giemsa-stained parasites [Fig 3C]. At 20 minutes after removal of the egress inhibitor, parasites lacking PV6 have the same surface area as wildtype parasites, but their surface area failed to increase by the two-hour time point [Fig 3C]. The volume of wildtype and mutant parasites also increased over time; interestingly, the volume of mutant parasites increased more than that of wildtype parasites [Fig 3D].

Similar results were obtained in a separate SBF-SEM experiment that used parasites that had been synchronized at the start of the first infection cycle but were not synchronized before entry into the second infection cycle [Supp Fig 1A]. In this experiment, wildtype parasites also assumed amoeboid shapes, whereas parasites lacking PV6 were spherical and had a smaller surface area than wildtype parasites [Supp Figs 4B and 4C]. Overall, these results show that the formation of the amoeboid form requires a PV6-mediated increase of the parasite surface area.

Interestingly, the surface area of merozoites was significantly smaller than that of intraerythocytic parasites observed 20 minutes after removal of the egress inhibitor, indicating that the parasites undergo a rapid expansion between release from the schizont and completion of invasion [Fig 3C]. The transformation of merozoites to the amoeboid form therefore consists of at least two steps: a transformation of a merozoite to a recently invaded parasite, followed by an expansion of the surface area and transformation to the amoeboid form. This second step does not occur in the absence of PV6, indicating that this protein has an essential role in the progression of the development of the parasites to this stage.

### The nucleus is elongated in amoeboid forms

Close analysis of the individual sections of the SBF–SEM experiments revealed that the connection between the limbs of the parasite can be extraordinarily thin. In the consecutive SBF-SEM sections shown in Fig 4A, three limbs of the parasite are visible, with a connection detected between the central limb and the other two limbs in only two 50 nm sections, with the connection barely perceptable in one of those two sections. In this parasite the connections between the limbs is thus likely less than 100 nm wide. Interestingly, the nucleus in this parasite spanned one of these very narrow connections, suggesting that the nucleus is flexible and can span multiple limbs. Although uncommon, such narrow connections between limbs were not unusual, although not all of them contained the nucleus [see additional examples in Supp Fig 5A].

**Figure 5.**
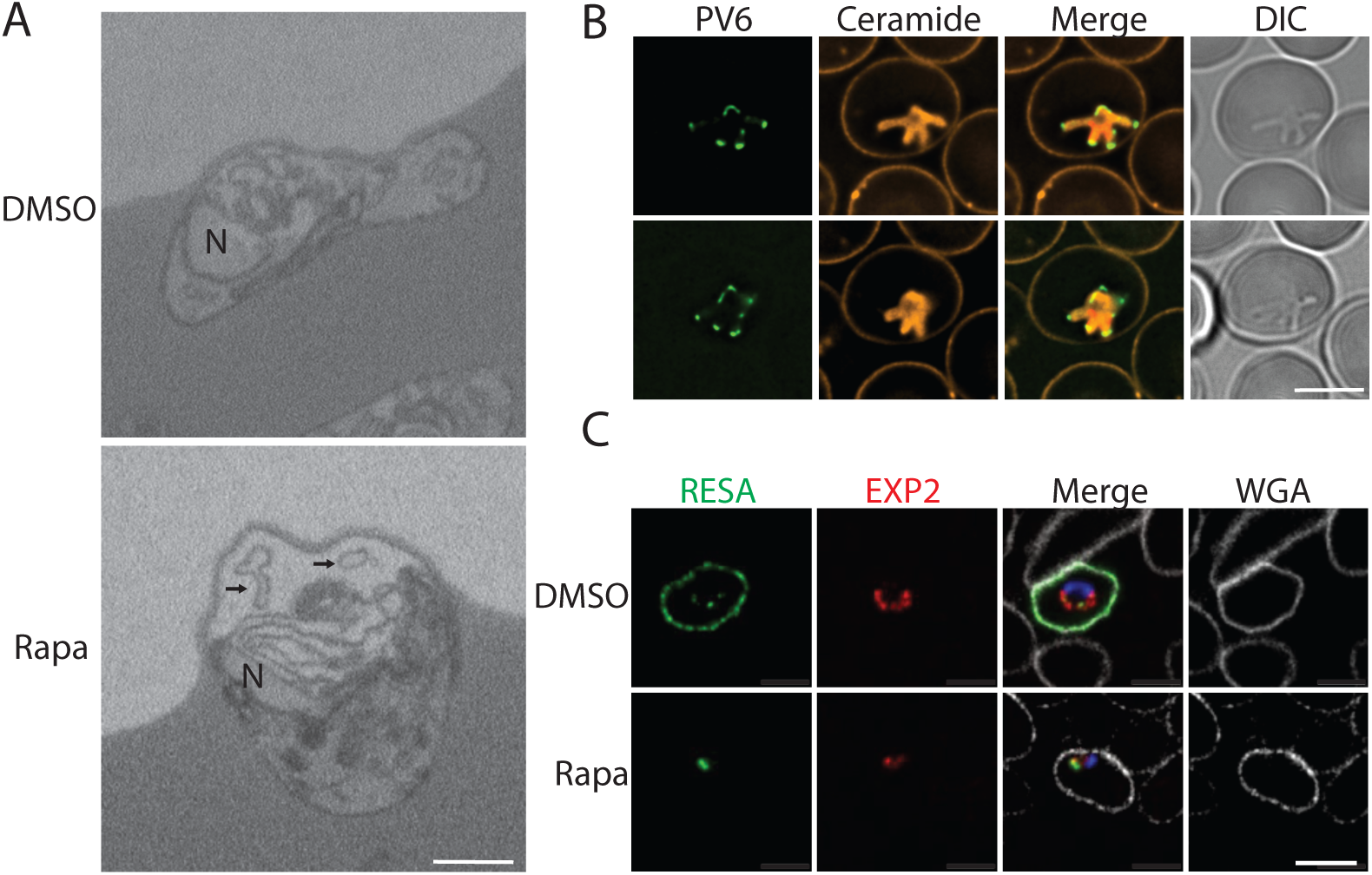
**Phenotype of *P. falciparum* parasites lacking PV6**. **A**. SBF-SEM sections from DMSO-treated (upper panel) and rapamycin-treated (lower panel) PV6 DiCre parasite culture, showing the accumulation of membrane whorls around parasites lacking PV6. Arrows indicate some of the lipid whorls. See Supp Fig 6 for additional images. **B.** Live-cell fluorescence imaging of parasites expressing an mNeonGreen-HA_3_-PV6 fusion (PfBLD717) one hour after removal of ML10. Membranes were labeled using C5-Bodipy-ceramide (orange). **C.** Immunofluorescence staining of DMSO-treated and rapamycin-treated PV6-DiCre parasites 6 hours after removal of ML10 using anti-RESA (green) and anti-EXP2 (red) antibodies. DNA was stained with Hoechst 33342 (blue) and the erythrocyte plasma membrane was visualized with Wheat Germ Agglutinin (WGA)-Alexa Fluor 647 (white). In all panels, the scale bar represents 5 µm.

Modeling of the nucleus in several SBF-SEM parasite models confirmed that the nucleus can extend into multiple limbs and also form an elongated, curved structure that fills one limb [Fig 4B, Supp Fig 5B], as also seen in previous 3D reconstructions^13,14^. Fluorescence microscopy of live parasites stained with a DNA dye two hours after the removal of egress inhibitor similarly revealed elongated nuclei that protrude into multiple limbs [Fig 4C, Supp Fig 5C]. Even in rounder parasites, the nucleus was extended or appeared to be circular, rather than spherical [Supp Fig 5C, right-hand panels]. This change in nuclear morphology likely starts after invasion has been completed, as the nucleus of invading merozoites has been found to form a barrier during invasion, slowing down the invasion process when it passes the tight junction^26^. Supporting this, in some amoeboid parasites, the nucleus was still spherical and formed the thickest region of the parasite [Supp Fig 5D, also see the DMSO-treated parasite in Fig 4B], suggesting that formation of the amoeboid form can occurs prior to nuclear remodelling. However, the nucleus in parasites lacking PV6 was more spherical than in wildtype parasites, indicating that amoeboid formation may be required for nuclear remodelling [Supp Fig 5A]. Similar changes to nuclear morphology have been detected in neutrophils, which undergo extreme shape changes as they cross the endothelium^27^. As is the case in those cells, the change in the nuclear morphology in parasites likely promotes the deformability of the parasite.

### Membrane ‘whorls’ accumulate around parasites lacking PV6

In SBF-SEM sections, parasites lacking PV6 were often detected close to the edge of the infected erythrocyte and in many cases whorls of what appeared to be lipid (membrane) material were detected in the space between the parasite and the parasitophorous vacuole membrane (PVM) [Fig 5A; see Supp Fig 6 for additional images]. As PV6 is a phospholipid transfer protein, the smaller surface area of the mutant parasites (and hence the PVM) and the accumulation of membrane whorls in the parasitophorous vacuole are consistent with the hypothesis that PV6 has a role in the transfer of lipids to the PVM to support the rapid expansion of parasites^16,18,28^.

**Figure 6.**
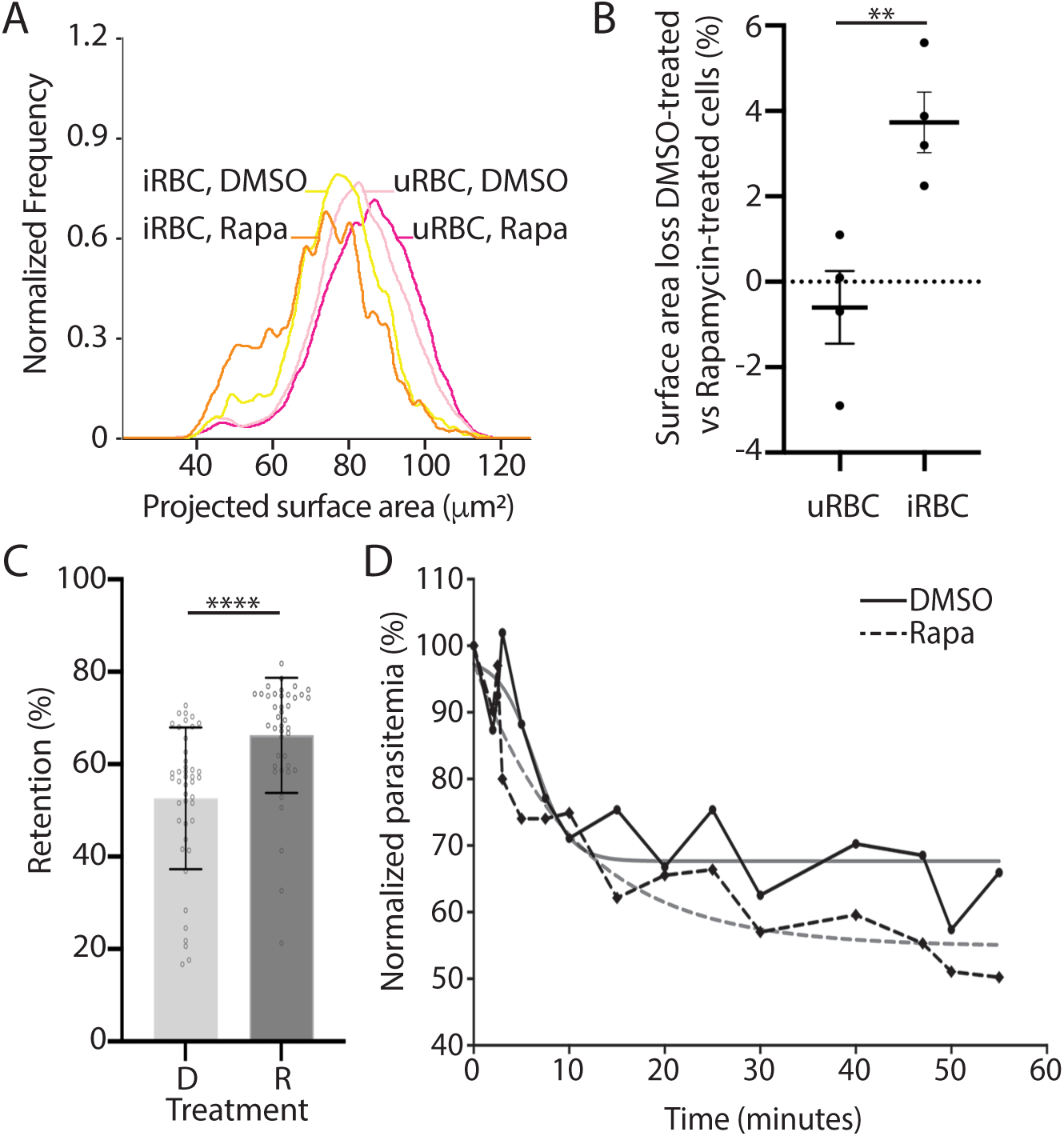
**Size and functional alteration of erythrocytes infected with parasites lacking PV6**. **A.** Surface area of uninfected erythrocytes (uRBCs) and erythrocytes infected with PV6-diCre parasites (iRBCs) 30 minutes after removal of egress inhibitor. The surface area of uRBCs and iRBCs in culture of wildtype (DMSO-treated) parasites and parasites lacking PV6 (Rapamycin-treated) was determined using imaging flow cytometry. Data represents four biological replicates. (T-test; **P˂0.01, Errors bars +/-SEM). **B.** Quantification of the surface area loss of uninfected (uRBC) and infected (iRBC) erythrocytes from cultures of DMSO-treated and rapamycin-treated parasites thirty minutes after removal of ML10, representing four experiments. **C.** Retention of erythrocytes infected with wildtype (DMSO-treated – D) parasites and parasites lacking PV6 (Rapamycin-treated – R) on a microsphiltration column. Data represents six biological replicates with a minimum of 6 technical replicates. Significance was determined using Mann-Whitney test; ****P˂0.0001, Errors bars represent SD. **D.** Retention of erythrocytes infected with PV6-DiCre parasites treated with DMSO or rapamycin in an *ex vivo* human spleen. Blood containing erythrocytes infected with DMSO-treated (solid black line) and rapamycin-treated parasites (dashed black line) approximately 6 hours after the removal of ML10 was continuously perfused through the spleen and the parasitemia was determined at the indicated times. Interpolation curves are shown in grey lines (R^2^=0.84 for DMSO and R^2^=0.90 for Rapa samples).

### PV6 activity impacts the remodelling of red blood cell following parasite invasion

To determine whether PV6 is in the correct subcellular location to distribute lipids to the PVM, the endogenous *pv6* gene was replaced with a form encoding a PV6-mNeonGreen-HA_3_ fusion protein [Supp Fig 7]. These modified parasites grew at a similar rate to wildtype parasites, indicating that the modification has no effect on the activity of the protein; a similar modification with an HA_3_ tag at the same position in the protein had no effect on parasite growth^17^. The fusion protein was detected at the tips of the limbs of the amoeboid forms, indicative of a localization in the PV, supporting previous findings using anti-PV6 antibodies [Fig 5B]^17^.

Next, we aimed to determine whether parasites lacking PV6 modify the host cell similarly to wildtype parasites. Six hours after the removal of egress inhibitor, as expected, the exported protein RESA was readily detected at the periphery of erythrocytes infected with wildtype parasites ^29^, however, no export of RESA was detected in erythrocytes infected with parasites lacking PV6, indicating that little to no export occurs in the absence of PV6 [Fig 5C]. Interestingly, the PVM protein EXP2^30^, which mediates protein export^31–33^, appeared to have a more restricted internal localisation in the PV containing parasites lacking PV6 (compared to a more peripheral localisation in the PV of wildtype parasites), indicating that EXP2 may not be in the correct location to export proteins [Fig 5C]. Interestingly, Riglar *et al.* showed that protein export does not appear to commence until about ten minutes after invasion^15^. Hence, parasites need to undergo a particular development after invasion to become competent to export proteins and PV6 seems to be required to complete this development. Alternatively, PV6 may have a role in export itself, but no evidence for such a role has been published.

It has been shown previously that infected erythrocytes have a smaller surface area than uninfected erythrocytes, as some of the erythrocyte membrane is internalized to form the PVM^9,34^. Using imaging flow cytometry, we also observed this phenomenon [Fig 6A].This analysis also revealed that the surface area of erythrocytes infected with mutant parasites is ∼4% smaller than erythrocytes infected with wildtype parasites [Fig 6B]. This difference is larger than the difference between the uninfected erythrocytes in the DMSO-treated and rapamycin-treated samples [Fig 6B]. Hence, PV6 activity mitigates the loss of surface area of infected erythrocytes. As smaller erythrocytes are more likely to be removed in the spleen^9,35^, PV6 thus may have an important role in preventing the removal of infected erythrocytes in the spleen by decreasing the loss of surface area.

### Absence of PV6 decreases passage of infected erythrocytes through the spleen

To determine whether the phenotypic changes of parasites lacking PV6 and their host cells affect the ability of the parasite to escape clearance by the spleen, we applied erythrocytes infected with wildtype parasites and parasites lacking PV6 to a microsphiltration column, which mimics filtration of erythrocytes through the IES *in vitro*^36,37^. Erythrocytes containing parasites lacking PV6 were retained in the column at a significantly higher rate than erythrocytes containing wildtype parasites [Fig 6C], suggesting that erythrocytes containing mutant parasites are more likely to be removed from the circulation when crossing the spleen filtration barrier. We tested this directly by perfusing erythrocytes infected with either wildtype parasites or parasites lacking PV6 through an *ex vivo* human spleen and found that erythrocytes infected with mutant parasites were cleared faster than the erythrocytes infected with wildtype parasites [Fig 6D, Supp Fig 8]. These results indicate that the modifications in the infected erythrocyte owing to the activity of PV6 play an important role in the survival of the parasite in the human host, and to our knowledge, this makes PV6 the first parasite protein that has been shown to promote the passage of infected erythrocytes through the host’s spleen.

## DISCUSSION

Together, our results reveal that the establishment of the malaria parasite *Plasmodium falciparum* in the host erythrocyte is a much more complex process than previously thought. Rather than passively adopting a ring shape after invasion, starting soon after (and possibily even during) the invasion process, the parasite undergoes a drastic metamorphosis, changing from a small, compact cell to a large, extended and flexible cell. All components required for this transformation are present in the merozoite and the host cell, as the transformation also takes place in the absence of nutrients or protein synthesis. In the approximately seventeen minutes required for the metamorphosis to occur, the cell is likely radically remodeled, such that the amoeboid-shaped parasite has little resemblance to the invading merozoite. Adopting an extended amoeboid form increases the surface area-to-volume ratio, which allows the parasite to be much more flexible than the spherical merozoite. This flexibility likely promotes passage of infected erythrocytes through the spleen and the microvasculature. Similarly, our data also reveal that the nucleus becomes extended and in some parasites is distributed over several limbs of the amoeboid, sometimes spanning connections between limbs less than 100 nm wide. This extended shape of the nucleus further likely adds to the flexibility of the cell as a whole and is a distinct change from the merozoite, which appears to have a rigid nucleus^26^. As gene expression is in part dictated by the 3D organization of the genome^38,39^, this distribution of the genome among the different lobes of the nucleus may impact the transcriptome of the parasite.

Although the mechanism of the merozoite-to-amoeboid metamorphosis remains unknown, it is of interest that this metamorphosis consists of at least two steps, starting with an expansion of the parasite between the merozoite stage and the recently invaded parasite, followed by a second expansion that likely involves the formation of the amoeboid form and requires the parasite phospholipid transfer protein PV6. Our finding that parasites lacking this protein are smaller and more spherical and have a smaller PVM may indicate that this protein has a role in the expansion of the PVM. These findings build upon the results presented by Dans *et al.*, who used lattice light-sheet microscopy to show that inhibiting PV6 activity also results in more spherical parasites^18^. As this protein is present in the PV after invasion, it is well positioned to transfer parasite-derived phospholipids, either phospholipids from the parasite plasma membrane or phospholipids secreted from the rhoptries^40,41^, to the PVM. This is consistent with a model in which the PVM is initially produced using lipids from the erythrocyte plasma membrane, and thus can form without PV6, and is subsequently expanded as the parasite increases in size during the transition to the amoeboid form, using parasite-derived phospholipids^34,42^. The results presented here indicate that phospholipids are also transferred to the surface of the infected erythrocyte, as erythrocytes containing parasites lacking PV6 have a smaller surface area than erythrocytes infected with wildtype parasites. As erythrocyte size is an important determinant of passage through the spleen^9^, the transfer of parasite phospholipids to the erythrocyte surface likely increases the survival of the parasite in the host, as indicated by the increased clearance of erythrocytes containing parasites lacking PV6 in the spleen. However, the mechanism for this phospholipid transfer remains obscure, as there is no known connection between the PVM and the plasma membrane of the erythrocyte and PV6 is confined to the PV^17^.

The transformation of the merozoite to amoeboid is a complex process that occurs in at least two distinct steps. PV6 is required for the parasite to undergo the second expansion and failure to progress leads to an inability of the parasite to export proteins. Previous results by Riglar et al. showed that export does not commence immediately after invasion; protein export was detected only approximately ten minutes after invasion^15^. Potentially, the second step of the transformation to the amoeboid form is required for the formation of the PTEX export system to be activated or positioned in the PVM to allow for export.

Although the complexity of the metamorphosis of the parasite is readily apparent from the large changes in the shape and surface of the cell, this complexity is further underscored by the possible functions of the few additional proteins that have been identified to have a role in this process: a PV protein (PV6), an inner membrane complex protein (IMC1g), an unidentified PM V target and a potential signalling protein (the phosphatase PP7)^17,25,43^. The metamorphosis from merozoite to amoeboid is an essential process, as parasites lacking PV6, IMC1g or PP7 do not develop, and in the case of IMC1g and PP7, die rapidly during culture *in vitro*. Our finding that the metamorphosis is driven by parasite proteins already in the merozoite indicates that this process could represent a target for intervention strategies. As the host is more likely to clear erythrocytes that contain parasites lacking PV6, this protein may be a more promising drug target than previously thought^18^.

Together, our results show that *Plasmodium falciparum* parasites undergo a rapid metamorphosis after invasion and that the alterations of the host cell that accompany this transformation increase the survival of the parasite inside the host.

## ACKNOWLEDGMENTS

The work in this study was supported in part by a Career Development Award from the Medical Research Council to CvO (MR/R008485/1), by funding from University of Paris City and Institut National de la Santé Et de la Recherche Médicale (INSERM) to CR, PAB and PAN and by funding to MRGR, MJB and LMC from the Francis Crick Institute (https://www.crick.ac.uk/) which receives its core funding from Cancer Research UK (CC2129), the UK Medical Research Council (CC2129) and the Wellcome Trust (CC2129).

The authors thank James Thomas (LSHTM) for advice on live imaging of parasite invasion, Don van Schalkwyk (LSHTM) for advice on live-dead staining of parasites, Juliana Chung (Université Paris Cité and Université des Antilles) for assistance with parasite culture and Paul Gilson (Burnet Institute) for sharing the EXP2 antibody. The authors acknowledge the facilities and the scientific and technical assistance of the LSHTM Wolfson Cell Biology Facility, with specific thanks to Liz McCarthy and Chris Chiu.

## ETHICS STATEMENT

The experiments using the *ex vivo* human spleen are part of the “SPLEENVIVO PROTOCOL” registered by the Comité de Protection des Personnes” (CPP) Ile de France and approved by the IRB with the folllowing number: 2015-02-05 MS2 DC. For the spleen used in this study, the subject signed a consent form prior to spleen retrieval and use for experiments.

## CONTRIBUTIONS

Conceptualization: AF, PAB, CvO

Methodology: AF, MR, MH

Investigation: AF, FML, ABD, NK, AF, AS, VC, CR, PAN, MR, CvO

Visualization: AF, CvO

Funding acquisition: MJB, LMC, PAB, SV, CvO

Project administration: CvO

Supervision: LMC, PAN, CvO

Writing – original draft: AF, CvO

Writing – review & editing: AF, FML,CR, MH, MJB, PAB, PAN, CvO

## MATERIALS AND METHODS

### Parasite culture, parasite strains and rapamycin treatment

*P. falciparum* parasites were cultured in human erythrocytes (sourced from UK National Blood Transfusion Service and Cambridge Bioscience, UK) at 3% haematocrit maintained in RPMI-1640 (Life Technologies) supplemented with 2.3 g/L sodium bicarbonate, 4 g/L dextrose, 5.957 g/L HEPES, 50 µM hypoxanthine, 0.5% AlbuMax type II (Gibco), and 2 mM L-glutamine (cRPMI) and incubated at 37°C in the presence of 5% CO_2_. *P. knowlesi* parasites were cultured in cRPMI further supplemented with 10% horse serum and was incubated in an atmosphere of 96% N_2_, 1% O_2_ and 3% CO_2_ ^22^

For routine culture, *P. falciparum* parasites were synchronized by isolating late-stage parasites on a Percoll cushion^44^, allowing the isolated parasites to invade fresh erythrocytes for 1-3 h and removing the remaining late-stage parasites on a second Percoll cushion. In the case of *P. falciparum*, late-stage parasites present in the culture were removed by treatment with 5% (w/v) sorbitol^45^, leaving only recently invaded parasites. To allow invasion to take place in a shorter window, parasites were treated with 25 nM ML10 (from a 100 µM stock) or 1 µM Compound 2 (from a 1 mM stock) to prevent egress^20,22^. When most schizonts appeared arrested, the parasites were pelleted, suspended in cRPMI containing erythrocytes at a hematocrit of 3% and incubated at 37°C unless otherwise stated.

All experiments were performed with *P. falciparum* strains 3D7, PfBLD529 clone A8^16^ and PfBLD529-466. The two different PfBLD529 parasite lines contain the identical floxed *pv6* locus (*pfa0210c*/PF3D7_0104200) but differ in the location of the diCre cassette. In PfBLD529 clone A8 this is present at the *sera5* locus^46^ whereas in PfBLD529-466 it is inserted in the *pfs47* gene^47^. As previously described, the native *pv6* gene is excised upon treating PfBLD529 parasites with rapamycin^16^.

Rapamycin treatment was performed on trophozoite-stage parasite at approximately ∼30 hours post-invasion. Parasites were incubated with 10 nM rapamycin (from a 100 µM stock solution in DMSO) at 37°C for one hour and controls were treated with an identical volume of DMSO^46,47^.

### Plasmids and transfection

Parasite line PfBLD529-466 was produced by transfecting 3D7 parasites with repair plasmid pBLD466 (also described as pBSPfs47DiCre)^47^. The resulting parasite line expresses diCre from the *pfs47* locus. This parasite line was subsequently transfected with plasmid pBLD529^16^ to produce PfBLD529-466. Parasite line PfBLD717 (producing the PV6-mNeonGreen fusion) was produced by amplifying the gene encoding mNeonGreen using primers CVO696b (CGCAGTCAATCTCGAGTTAGTAAAGGAGAAGAAGATAATATGGCAAG) and CVO697b (CGTAAGGGTACTCGATTTATACAATTCATCCATTCCCATAACATCTGTAAATG). The resulting DNA fragment was cloned using InFusion (Takara Bio) into pBLD708 ^17^ that had been digested with AvrII and SpeI, producing plasmid pBLD717. Transfections were performed using the schizont transfection protocol as described previously ^46^. Briefly, schizonts purified on a Percoll cushion were allowed to invade fresh erythrocytes for 1-3 hours. The residual schizonts were isolated on a Percoll cushion, pelleted, washed with cRPMI and suspended in a solution consisting of 100 µL AMAXA nucleofection P3 Primary Cell transfection reagent and 10 µL TE containing 15–30 µg targeting plasmid and 20 µg plasmid pDC2-Cas9-hDHFRyFCU ^17,47^ encoding Cas9 and the appropriate guide RNA. Transfection was carried out using the AMAXA nucleofection device (Lonza) using program FP158. Immediately following transfection, schizonts were mixed with fresh erythrocytes and 3 ml cRPMI and incubated with shaking at 37°C for 30 minutes to facilitate invasion. Drug selection with 2.5 nM WR99210 was started after 24 hours and sustained for 7 days. Transfected parasites were typically recovered 3–4 weeks after transfection.

### Analysis of parasite development using Giemsa-stained samples

To analyse and measure the development of parasites, PfBLD529-466 (PV6 diCre) parasites were tightly synchronized using a Percoll-Sorbitol strategy and then treated with rapamycin and ML10 as outlined above. At the indicated time point after invasion, the parasites were pelleted, smeared on a microscope slide, fixed with methanol and stained with Giemsa. The parasites were imaged using an Olympus BX51 microscope equipped with an Olympus SC30 camera and a 100x oil objective, controlled by CellSens software. Analysis of the images was performed using FIJI. The contour of each visible parasite was delineated using the FIJI freehand selection tool and the area was determined utilizing the area measurement function. Each experiment was performed in triplicate with a minimum of three biological replicates. Graphpad Prism v10 was used for statistical analysis.

Analysis of the ability of the parasite to form amoeboids was first performed using live microscopy. Parasites were tightly synchronized using the Percoll-reinvasion-Sorbitol strategy strategy detailed above. The parasites were treated with rapamycin at 30 hours post-invasion and subsequently arrested at the late schizont stage with ML10 to inhibit egress ^22^. When appropriate, the initiation of synchronized invasion was triggered by the removal of ML10. At the indicated time point after removal of ML10, the parasites were collected for analysis. Prior to imaging, the cells were labelled with 5 µg/mL Hoechst 33342 and allowed to settle in a 6-well Ibidi µ-slide with a glass bottom. Parasites were observed using a 100X oil immersion objective on a Nikon Ti-E inverted microscope equipped with a Hamamatsu ORCA-Flash 4.0 Camera and Piezo stage driven by NIS elements version 5.3 software. Image J was employed for image processing, with adjustments made to brightness and contrast using the reset command for autoscale. Images were sized in Photoshop and figures were produced using Illustrator. To measure the area and circularity of the parasites, the contour of each visible parasite was delineated using the FIJI freehand selection tool followed by additional measurements using the area and circularity functions. Circularity values range from 0 to 1 (derived from the area and perimeter of the cell). Higher values indicate shapes of increasing circularity, with a perfect circle characterised by a value of 1. Each experiment was performed in triplicate with a minimum of three biological replicates. Graphpad Prism v10 was used for statistical analysis.

To visualize parasites using fluorescent ceramide, the culture was incubated at 37°C in the presence of 7 mM Bodipy C5-ceramide in complex with BSA (diluted from a 100 mM stock in PBS) and 2 µM 5-SiR-Hoechst^48^ in RPMI without Albumax for a minimum of 20 minutes. The culture was pelleted at 2000 x g, suspended in RPMI without Albumax and diluted to a hematocrit of 0.6% using RPMI without Albumax before loading into a poly-L-lysine-coated 6-well Ibidi µ-slide. The parasites were visualized on a Nikon Ti-E inverted microscope as described above.

To determine the effect of the protein synthesis inhibitor cycloheximide or Hanks’ Balanced Salt Solution containing Mg^2+^ and Ca^2+^ (HBSS) on amoeboid formation, synchronized late-stage parasites were purified on a Percoll cushion, washed with, and then suspended in, cRPMI and incubated in the presence of ML10 to prevent egress. When most parasites appeared arrested, 5 µM cycloheximide or an equivalent volume of DMSO was added and the parasites were incubated for an additional hour. The parasites were then pelleted, mixed with erythrocytes, in the presence of either DMSO or cycloheximide. Half of the DMSO-treated parasites was removed, pelleted, washed four times with HBSS and suspended in HBSS and all samples were incubated at 37°C to allow invasion to take place. After two hours, the parasites were diluted to a hematocrit of 0.6% in RPMI before being loaded into a poly-L-lysine-coated 6-well Ibidi µ-slide. The parasites were visualized live on a Nikon Ti-E inverted microscope as described above.

To determine the timing of amoeboid formation, late-stage parasites were purified on a Percoll cushion and incubated at 37°C in the presence of 10 nM ML10 to prevent egress. When most parasites appeared arrested, the parasites were pelleted and mixed with erythrocytes to allow invasion to occur. At various time points starting at fifteen minutes, 200 µl of culture was removed, pelleted and fixed with 2% paraformaldehyde, 3% glutaraldehyde in 0.1 M phosphate buffer (Karnovsky’s fixative). After a minimum of one hour of fixation, the cells were pelleted and suspended in 1 ml PBS to adjust the hematocrit to 0.6%. The cells were subsequently viewed using poly-L-lysine-coated 6-well Ibidi µ-slides as described above using a Nikon Ti-E inverted microscope.

### Video microscopy of invasion and amoeboid formation

Parasites were tightly synchronized using the Percoll-reinvasion-Sorbitol strategy described above and subsequently treated with ML10 to inhibit egress. Late-stage parasites were isolated on a Percoll cushion, suspended in 30 ml cRPMI containing 25 nM ML10 and incubated at 37°C. When most parasites appeared arrested, 5 ml of the purified schizont suspension was pelleted and resuspended in warm cRPMI containing erythrocytes at a hematocrit of 3%. The parasitemia was adjusted to approximately 10% and the hematocrit was adjusted to 0.6% using warm cRPMI. 200 μl was loaded into a poly-L-lysine coated μ-Slide VI 0.4 (Ibidi) contained in a small warm bead bath. The slides were transferred to a Nikon Ti-E inverted microscope housed in a pre-warmed 37 °C chamber with an atmosphere of 5% CO_2_. Egress, invasion of the erythrocyte by the released merozoites and the establishment of the parasites in the erythrocyte were imaged using a 100× oil immersion objective and an ORCA-Flash 4.0 CMOS camera (Hamamatsu), at a rate of 1 frame/sec for one hour. Videos were acquired and processed using Nikon NIS-Elements software.

### Immunofluorescence microscopy

Schizonts, either treated with DMSO or rapamycin as described above and subsequently purified on a Percoll cushion, were allowed to invade erythrocytes for 1-6 hours and were then fixed in a solution containing 8% paraformaldehyde, 0.01% glutaraldehyde in PBS for 1 hour at room temperature^49^. The fixed parasites were permeabilized with 0.1% Triton X-100 in PBS for 10 minutes at room temperature, blocked with 3% BSA in PBS overnight at 4°C and then stained with mouse anti-RESA antibodies (1:500, obtained from the Antibody Facility at The Walter and Eliza Hall Institute of Medical Research) and rabbit anti-EXP2 antibodies (1:1000, a kind gift of Paul Gilson) overnight at 4°C. parasites were washed three times with PBS and incubated at room temperature in the presence of fluorescently labelled secondary antibodies (1:10,000; labels used were Alexa Fluor 488 and Alexa Fluor 568), wheat germ agglutinin (WGA)-Alexa Fluor 647 (2 µg/mL) and Hoechst 33342 (5 µg/mL) for an additional 30 minutes. Following three washes with PBS, 1.5 µL of parasites suspension was placed on a polyethyleneimine-coated glass slide, mixed with 1.5 µL of Vectashield anti-fade mounting medium, and covered with a cover glass sealed with nail polish. Parasites were imaged on a Zeiss LSM 880 confocal microscope controlled by Zen Black version 2.3 software. The emission wavelengths used were 462 nm (Hoechst), 552 nm (Alexa 488), 633 nm (Alexa 568) and 693 nm (WGA-Alexa Fluor 647). FIJI software was used to separate, crop and adjust the brightness and contrast of the images.

### Serial block-face scanning electron microscopy (SBF-SEM)

To obtain 3D models of intraerythrocytic parasites Infected erythrocytes were prepared twice for SBF-SEM. In one experiment (Experiment 1) PV6-diCre parasites were synchronized and treated with rapamycin as described above. Compound 2 was then added (1 µM final concentration) to prevent egress. When most parasites appeared arrested, the parasites were pelleted and resuspended in warm cRPMI to remove Compound 2 and the culture was divided over four flasks. At the indicated time, a sample was removed from the cultures and immediately fixed in Karnovsky fixative and stored at 4°C. After 40 minutes, Compound 2 was added to the cultures to prevent further egress and keep the parasites in a narrow window of invasion. The cultures were subsequently pelleted at 600 xg and suspended in molten 1% low-melting point agarose in 0.1 M phosphate buffer, pH 7.4 and placed in a microcentrifuge tube. The samples were stored at 4°C and prepared for microscopy as follows. Fixed agarose blocks containing cultures were washed three times for 10 mins in 0.1M phosphate buffer pH 7.4. The samples were then osmicated for 2 hours in 2% aqueous osmium tetroxide. The samples were then washed three times for 10 minutes in distilled water and dehydrated by three 10-minute washes in 30%, 50%, 70%, 90%, and 100% ethanol. The final dehydration step involved 100% dry acetone (2 times 30 mins). The blocks were infiltrated overnight with 50% Durcapan resin in dry acetone before being transferred to 100% Durcapan resin for a further 12 hours after which time they were cured for 24 hours at 60°C. Pieces of resin blocks containing samples were trimmed and mounted onto aluminium pins using conductive epoxy glue and silver dag, and then sputter coated with a layer (10-13 nm) of gold in an Agar Auto Sputter Coater (Agar Scientific). Before SBF-SEM imaging, ultrathin sections microscope (JEOL) using a OneView 16-megapixel camera (Gatan-Ametek), to verify sample quality. Samples were then imaged in a Merlin VP compact high resolution scanning electron microscope (Zeiss) equipped with a 3View 2XP stage (Gatan-Ametek) and an OnPoint backscattered electron detector (Gatan-Ametek). Image series were acquired under high vacuum, using a focal charge compensation device (Zeiss) set to 100% output. The following imaging conditions were used: (1) 1.8kV, 20 µm aperture, 5 nm pixel size, 1-3 μs pixel time, 70 nm section thickness. SBF-SEM data were processed using the IMOD software package^50^. Briefly, image stacks were assembled, corrected (for z scaling and orientation) and aligned using eTOMO. Parasites and erythrocytes were segmented manually using 3dmod (part of the IMOD software package). Movies of data and models were produced using a combination of 3dmod and Fiji^51^. Volume, surface area and length measurements for different models were obtained in 3dmod. Statistical analysis of the data was performed using GraphPad Prism v10.

In the other experiment (Experiment 2) PV6-diCre parasites were synchronized and then treated with rapamycin approximately 30 h later. The parasites were harvested when most of them had progressed to the next cycle. The parasites were then fixed by adding 8% formaldehyde in 0.2 M Sorensen’s phosphate buffer (pH 7.4) in a 1:1 ratio with the growth medium, followed by fixation in 2.5% glutaraldehyde/4% formaldehyde in 0.1 M phosphate buffer for 30 min. Next the parasites were pelleted at 300 x g, embedded in 4% agarose, and cut into 1 mm^3^ blocks on ice, and washed in PB. The parasite-agarose blocks were then prepared using a modified NCMIR protocol^52^ by being post-fixed in 2% osmium tetroxide/1.5% potassium ferrocyanide for 1 h, incubated in 1% w/v thiocarbohydrazide for 20 min before a second staining with 2% osmium tetroxide for 30 min, then incubated overnight in 1% aqueous uranyl acetate at 4°C. The blocks were stained with Walton’s lead aspartate for 30 minutes at 60°C and dehydrated through an ethanol series on ice, incubated in propylene oxide, followed by overnight incubations in a 1:1 propylene oxide/Durcupan resin mixture and twice in Durcupan resin, and embedded in Durcupan resin according to the manufacturer’s instructions (TAAB Laboratories Equipment Ltd.). Images were collected using a 3View 2XP system (Gatan Inc.) mounted on a Sigma VP Scanning Electron Microscope (Carl Zeiss). Images were collected at 1.8 kV using the high-current setting, with a 20 μm aperture, at 8 Pa chamber pressure, 50 nm cutting thickness, and a pixel size of 2.5 × 2.5 nm with a 1 μs dwell time.

### Surface area analysis of infected erythrocytes using ImageStream

Parasites were prepared for ImageStream analysis by synchronizing parasite invasion using Percoll cushions and ML10 as described above. Parasites were washed and allowed to invade erythrocytes for 30 minutes and 90 minutes. The cells were pelleted at 2000 x g and suspended in 1% glutaraldehyde in PBS at a hematocrit of 2%. Subsequently, the cells were washed twice with PBS and then suspended in PBS containing 1% Albumax at a hematocrit of 1%. Imaging flow cytometry was performed using ImageStream X Mark II (AMNIS part of EMD Millipore) as described previously^53^. Briefly, infected erythrocytes were identified by labeling cells with SYBR Green for one hour followed by a wash with PBS. Images to determine RBC dimensions and morphology were obtained using brightfield images (60X magnification) and subsequently processed using IDEAS v6.2 software (AMNIS). Focused cells and single cells were respectively selected using the features gradient RMS and aspect ratio versus area. Front views were selected and analyzed by the mask “Object” and the features of circularity, perimeter and area. At least 6000 front views of focused single RBC were analyzed for each sample.

### Microsphiltration

Wildtype parasites and parasites lacking PV6 were allowed to invade erythrocytes for 4 hours in cRPMI and then washed and resuspended at 1% haematocrit in PBS containing 1% Albumax (Life Technologies). The retention rate was evaluated by microsphiltration (microsphere filtration) assay in 96 wells microplates as previously described^54^. The microsphiltration assay uses layers of metallic beads with different sizes to mimic the filtration of erythrocyte in the red pulp of the spleen^36^. The retention of infected erythrocytes was analysed by flow cytometry (BD FacsCantoII, BD Biosciences) comparing parasitemia of upstream versus downstream samples stained by Hoechst DNA dye. For each microplate, 95% of healthy blood donor (Etablissement Français du Sang, Lille, France) stained with carboxyfluorescein diacetate succinyl ester (CFSE) was mixed either with 5% of unstained healthy blood donor (RBC, negative control) or with 5% of 1% glutaraldehyde-fixed unstained healthy blood donor (rigid RBCs, positive control). The results from the experiment were analyzed when the retention rate <10% for negative control and >90% for positive control.

### Perfusion of *ex vivo* spleen

Parasites were synchronized and treated with DMSO or rapamycin as described above. One ml pellet of the DMSO-treated culture was stained with Cell Trace Violet according to manufacturer’s instructions and mixed with 1 ml of the rapamycin-treated culture that had been stained with stained with Cell Trace Far Red. The mixture was diluted with 18 mL of healthy erythrocytes. Parasites DNA was stained with SybrGreen for the evaluation of parasitemia. The 20 ml sample was then suspended at 10% hematocrit in Krebs-albumin solution. The healthy human spleen was obtained from a patient who underwent a distal splenopancreatectomy for pancreatic disease as described previously^55^. The splenic artery was cannulated through a catheter before being connected to the perfusion platform. The spleen was co-perfused with the mixed erythrocyte population for about 60 minutes through the spleen at 37°C. Samples were retrieved from a reservoir in the circuit at different time points and evaluated by flow cytometry analysis to determine the persistence of each subpopulation of infected erythrocytes. The parasitemia of each subpopulation was normalized at 100% from the start of perfusion and monitored during the time of perfusion. Owing to the scarcity of human spleens available for experimentation, this experiment was performed only once.

## SUPPLEMENTARY MOVIES

**Supplementary movie 1:** Live-cell imaging of *Plasmodium knowlesi*-infected erythrocyte. Note the transition from an amoeboid shape to a round shape, after which the parasite once again assumes an amoeboid shape. Image was captured in real time.

**Supplementary movie 2:** Live-cell imaging of *Plasmodium knowlesi*-infected erythrocyte. Note the transition from an amoeboid shape to a round shape, after which the parasite once again assumes an amoeboid shape. Image was captured in real time.

**Supplementary movie 3**: Serial block face scanning electron microscopy dataset of DMSO-treated parasites 2 hours post-removal of the egress inhibitor. The section thickness (Z resolution) is 50 nm, and each dataset consist of 439 images (pixel dimension and X-Y resolution). Note that this experiment was set up with a high ratio of schizonts to uninfected erythrocytes to maximize invasion. This resulted in a high number of free merozoites.

**Supplementary movie 4:** Serial block face scanning electron microscopy dataset of rapamycin-treated parasites 2 hours post-removal of the egress inhibitor. The section thickness (Z resolution) is 70 nm, and each dataset consist of 221 images (pixel dimension and X-Y resolution). Note that this experiment was set up with a high ratio of schizonts to uninfected erythrocytes to maximize invasion. This resulted in a high number of free merozoites.

**Supplementary movie 5:** SBF-SEM imaging and 3D model of erythrocytes infected with wildtype parasites or parasites lacking PV6. Movies consists of consecutive SBF-SEM sections. Erythrocytes and parasites used in the model are labeled in red and cyan, respectively.

## SUPPLEMENTARY METHODS

### Growth assay

To determine the parasite growth rate, the parasites were first tightly synchronized using a Percoll/Sorbitol strategy as described in the Methods section. Thirty hours post-invasion, the cultures were treated with rapamycin or DMSO and adjusted to a parasitaemia of 0.1%-0.2% and 2% haematocrit. Growth was assayed every two days for a total of six days by collecting a 50 µl aliquot which was fixed in an equal volume of fixative solution (8% paraformaldehyde, 0.2% glutaraldehyde in PBS) in a 96-well plate. When ready, the samples were prepared for analysis by flow cytometry: the fixative was removed and the cells were washed twice with PBS and suspended with 200 µl of PBS/SYBR Green 1 (1:5000, Life Technologies) for 30 minutes in the dark. Finally, the samples were diluted in PBS (1:5) and the parasitaemia was measured using an Attune cytometer (Thermofisher) using the following laser settings: forward scatter 125 V, side scatter 350V and blue laser (BL1) 530:30 280V. Each experiment was performed in triplicate with a minimum of three biological replicates. Graphpad Prism v10 was used for statistical analysis.

### Merozoite number

The quantification of merozoites per schizont was conducted using Giemsa-stained smear of mature parasite enriched through a Percoll cushion ^44^. The parasites were imaged using an Olympus BX51 microscope equipped with an Olympus SC30 camera and a 100x oil objective, controlled by cellSens software. Each experiment was conducted in triplicate with a minimum of three biological replicates. Statistical analysis was performed using GraphPad Prism v10.

### Parasite invasion rate

Parasite invasion assays were performed as described previously ^56^. Briefly, DMSO-treated and rapamycin-treated parasites were diluted to a parasitaemia of 1% and a haematocrit of 2%. At that point, a 50 µl starting sample (designated as H0) was collected and fixed in a PBS solution containing 8% paraformaldehyde, 0.01% glutaraldehyde. After a 24-hour incubation period, a second sample was collected and fixed (designated as H24). The parasites were labelled with SYBR Green I (1:5000, Life Technologies) and the parasitaemia was quantified using an Attune cytometer (Thermofisher; see detailed protocol above). Parasite invasion rate was determined as the ratio of H24/H0 parasitaemia. Each experiment was conducted in triplicate with a minimum of three biological replicates. Statistical analysis was performed using GraphPad Prism v10.

### Schizonts DNA content analysis

Highly synchronous schizonts were fixed and prepared as described above (“growth assay” section). SYBR Green fluorescence was analysed using an Attune cytometer (Thermofisher) using the following laser settings: forward scatter 125 V, side scatter 350V and blue laser (BL1) 530:30 280V. At least 100,000 cells were analysed per sample in three biological replicates. Data analysis was performed using FlowJo software.

### Live/dead staining

Parasites were treated with 10 nM rapamycin or an equivalent volume of DMSO for 30 minutes approximately 30 hours after invasion. The cultures were adjusted to a parasitaemia of 5% and a hematocrit of 3%. Parasites were collected at that point, then the next day (Schizonts) and at 1, 4, 8, 24 and 32 hours post-invasion. Upon collection, the samples were stained with 200 nM Mitotracker DeepRed (Thermo Fisher) for 15 minutes, washed with PBS and then fixed in an equal volume of fixative solution (8% paraformaldehyde, 0.2% glutaraldehyde in PBS) containing SYBR Green I (1:5000). After a minimum of 30 minutes, cells were washed with PBS and diluted in PBS (1:5) for cytometry. The analysis was performed on an Attune cytometer. Mitotracker-positive, SYBR Green-positive parasites were considered viable, Mitotracker-negative, SYBR Green-positive parasites were considered non-viable. The proportion of live parasite (stained with Mitotracker DeepRed) was compared to the proportion of all parasites (dead and viable) stained with SYBR Green as previously described ^57^. Two biological replicates were performed, each performed in triplicate. Data analysis was performed using FlowJo software.

### Immunoblotting

Parasites were isolated on a Percoll gradient, washed several times with RPMI without Albumax and subsequently suspended in SDS-PAGE buffer at a concentration of 10^5^ parasites/ul. Parasite lysates representing 6x10^5^ parasites were separated by SDS-PAGE, transferred to nitrocellulose using a BioRad Turboblot, blocked using PBST containing 5% milk and probed with rabbit anti-PV6 (1:1000)^16^ or anti-HA (3F11, 1:2000; Merck) Antibodies were visualized by incubating the blots with the appropriate HRP-linked anti-rabbit (PV6) or anti-rat (HA) secondary antibodies and developing using Clarity ECL Western blotting substrates (Bio Rad). Blots were imaged using a ChemiDoc (BioRad).

---

**Supplementary Figure 1:**
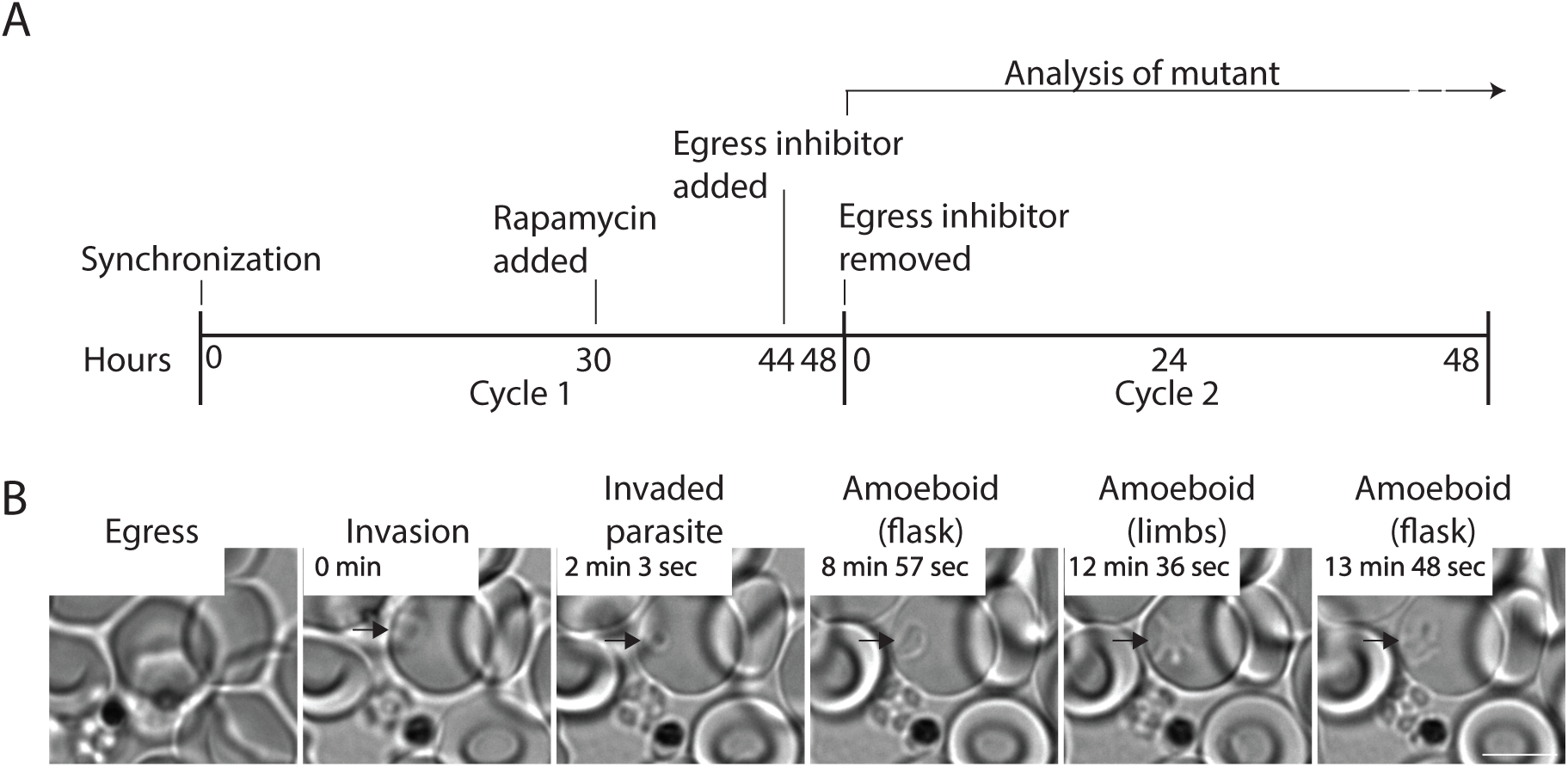
Phenotypic analyses of parasites lacking PV6. **A.** Schematic representation of the protocol used to analyse the phenotype of the *P. falciparum* PV6 DiCre parasites. Synchronized parasites were treated with a minimum of 10 nM rapamycin (or an equivalent volume of DMSO) for 1 hour at the trophozoite stage (∼30 hours post-invasion) in the first cycle. Close to the end of the first cycle, an egress inhibitor (either Compound 2 or ML10) was added to arrest the parasites at a very late schizont stage. When most of the parasites appeared arrested, the egress inhibitor was removed to initiate a round of synchronized invasion, starting cycle 2, allowing for observation of the phenotype of PV6 over time. **B.** Live-cell imaging of amoeboid formation. The frame rate was set at one frame/second, time starting from invasion is indicated in each panel. The arrows indicate the parasite. Scale bar represent 5 µm.

**Supplementary Figure 2.**
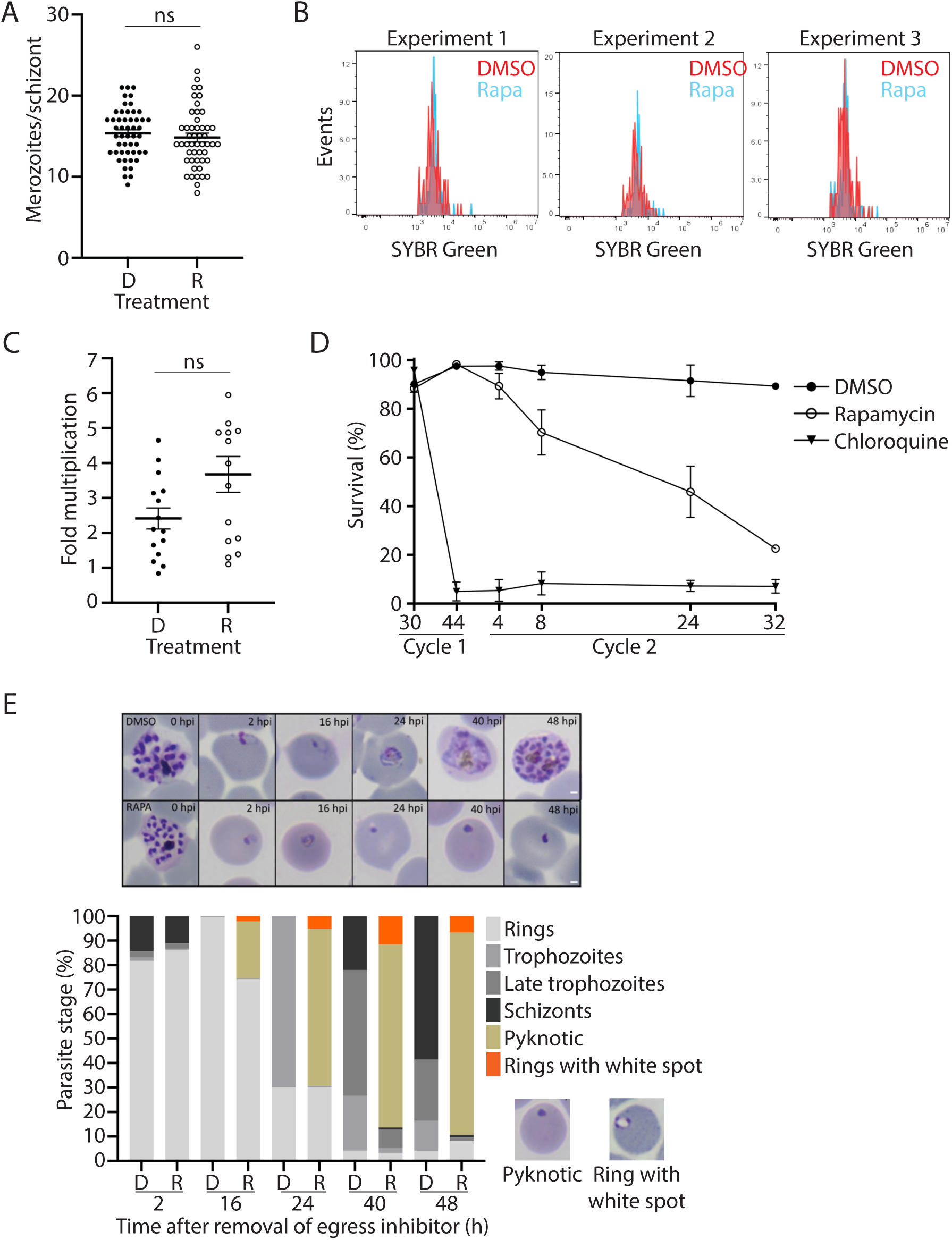
Phenotypic analyses of parasites lacking PV6. **A.** Number of merozoites per schizont in PV6-diCre (PfBLD529) parasites treated with DMSO (D) or rapamycin (R). Results are presented as the average number of merozoites per schizont of three biological replicates (a minimum of 15 schizonts were counted per replicate). Error bars indicate the error of mean. Mann–Whitney U test was performed for statistical analysis (ns: not significant). **B.** DNA content of the schizonts of DMSO-treated and rapamycin-treated PV6-diCre (PfBLD529) parasites. Shown are histograms from three individual experiments representing the DNA content (determined using SYBR Green staining and flow cytometry) of Percoll-purified PV6-diCre (PfBLD529) parasites treated with DMSO (red) or rapamycin (blue). At least 100,000 events were counted per sample (only events representing infected erythrocytes are shown). No obvious difference in the DNA content between the DMSO-treated and rapamycin-treated parasites was detected. **C.** Invasion rate of DMSO-treated and rapamycin-treated PV6-diCre (PfBLD529) parasites. Schizonts were incubated with fresh erythrocytes and the parasitemia was determined immediately (0 h) and after 24 hours (24 h). The invasion rate was determined as the ratio of the parasitaemia at the two time points (24 h/0 h). Five biological replicates, all in triplicate, were performed. Error bars indicate the error of mean. The Mann–Whitney U test was performed for statistical analysis (ns: not significant). **D.** Survival of PV6-diCre (PfBLD529) parasites treated with DMSO or rapamycin at the trophozoite stage. Parasite viability was determined by staining parasites with Mitotracker DeepRed and SYBR-Green. Invasion in the second cycle was synchronized using ML10. Viable (Mitotracker DeepRed-positive) parasites were compared to the percentage of all SYBR green-positive parasites (dead and viable parasites). Analysis was performed at the indicated time after removal of ML10. Chloroquine was used as control for detection of non-viable parasites. Results are presented as the mean (+/-SD) of two biological replicate assays performed in triplicate. E. Top panel: Giemsa-stained thin films showing the development of DMSO-treated (top row) and rapamycin-treated (bottom row) parasites at the indicated times after removal of the egress inhibitor. The scale bar represents 1 µm. Bottom panel: quantification of parasite developmental stages or phenotype of parasite development at the indicated time after removal of ML10. Rapamycin-treated parasites (R) formed rings but showed an accumulation of rings with white spots and pyknotic forms at an early stage (see examples of these phenotypes under the legend). These parasites did not develop to the trophozoite stage. Results are presented as the means of the count of two biological replicates.

**Supplementary Figure 3.**
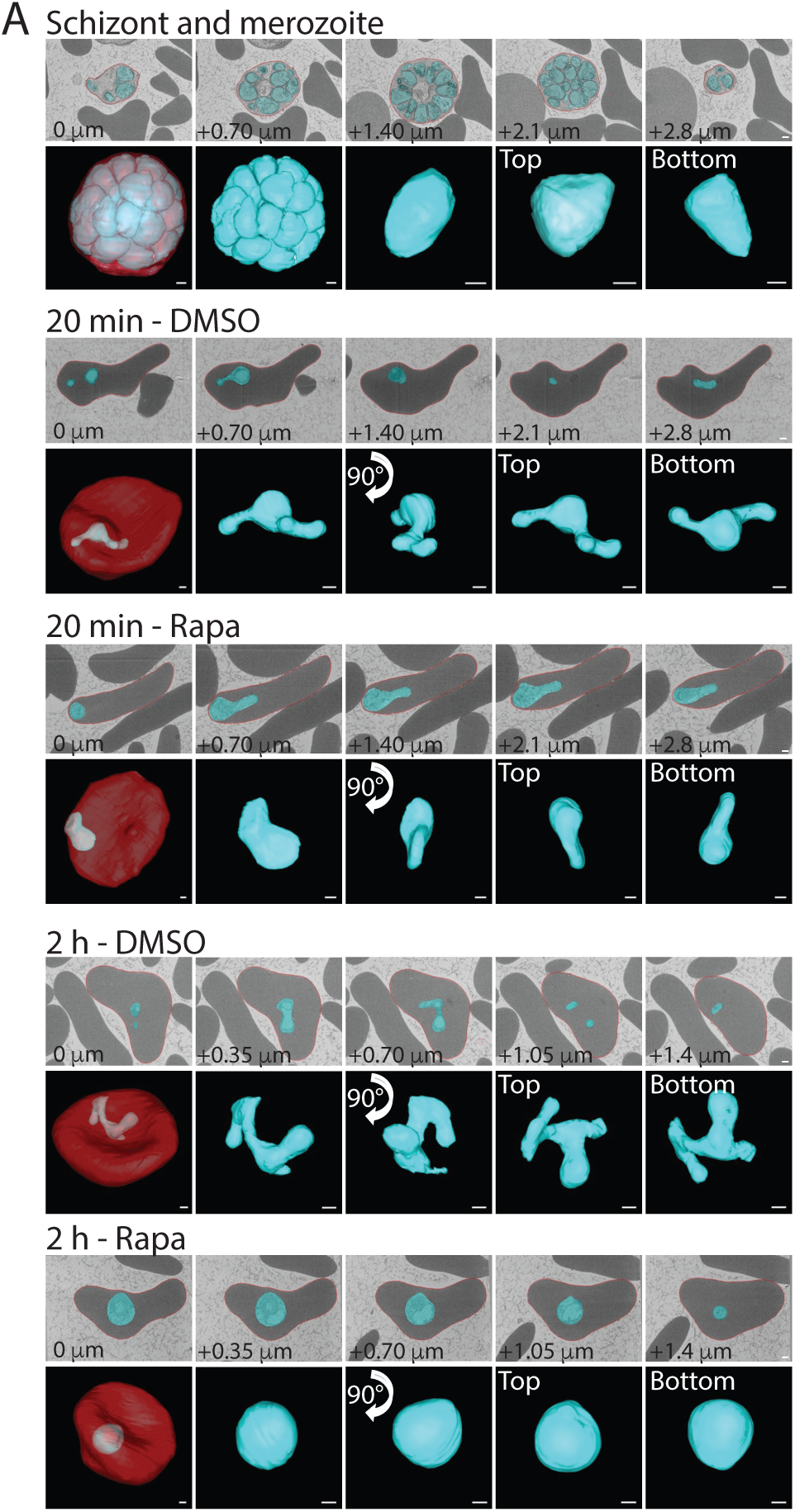
Additional three-dimensional views of the models presented in. Figure 2. Three-dimensional models from SBF-SEM data of the parasites presented in Figure 2, illustrating the shape of the parasite (cyan) and its positioning within the erythrocyte (red) and additional SEM-SBF sections. The scale bars represent 500 nm.

**Supplementary Figure 4.**
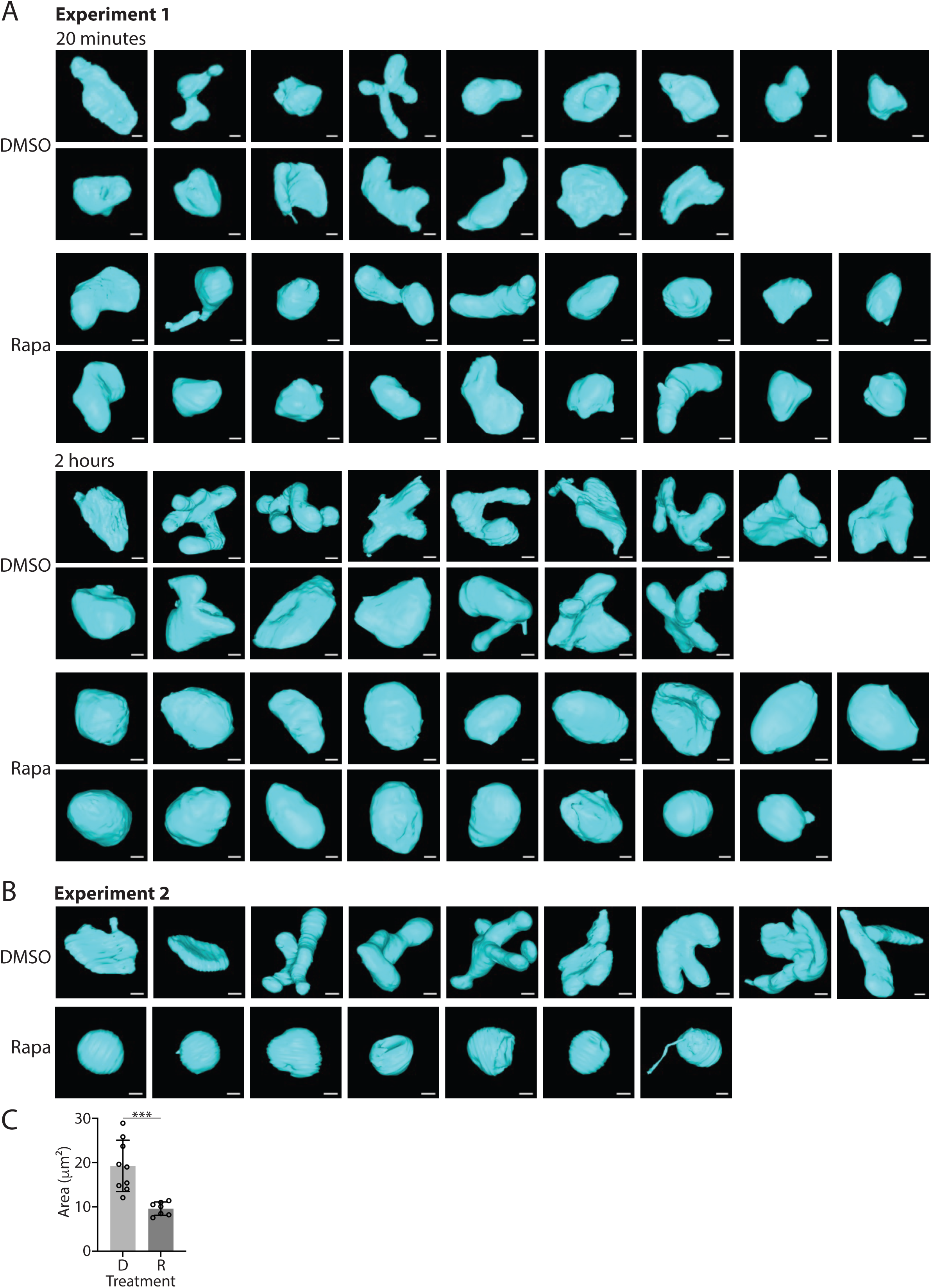
Three-dimensional models of PfBLD529 parasites treated with DMSO or rapamycin. **A.** These models were obtained in the same experiment as those presented in Figure 2 and Supp Figure 3. The invasion of these parasites was carefully synchronized using an egress inhibitor to allow the development of the parasites to be followed over time. Time indicated refers to the time after removal of egress inhibitor. The scale bars represent 500 nm. **B.** Three-dimensional models from SBF-SEM data of DMSO-treated and rapamycin-treated parasites for which invasion was not synchronized. The parasites had been synchronized at the start of cycle 1 [Supp Fig. 1] and were allowed to progress to the next cycle without further synchronization. The scale bars represent 500 nm. **C.** Surface area of the parasites shown in panel B; D-DMSO, R-rapamycin. The Mann–Whitney U test was performed for statistical analysis (***P < 0.001).

**Supplementary Figure 5:**
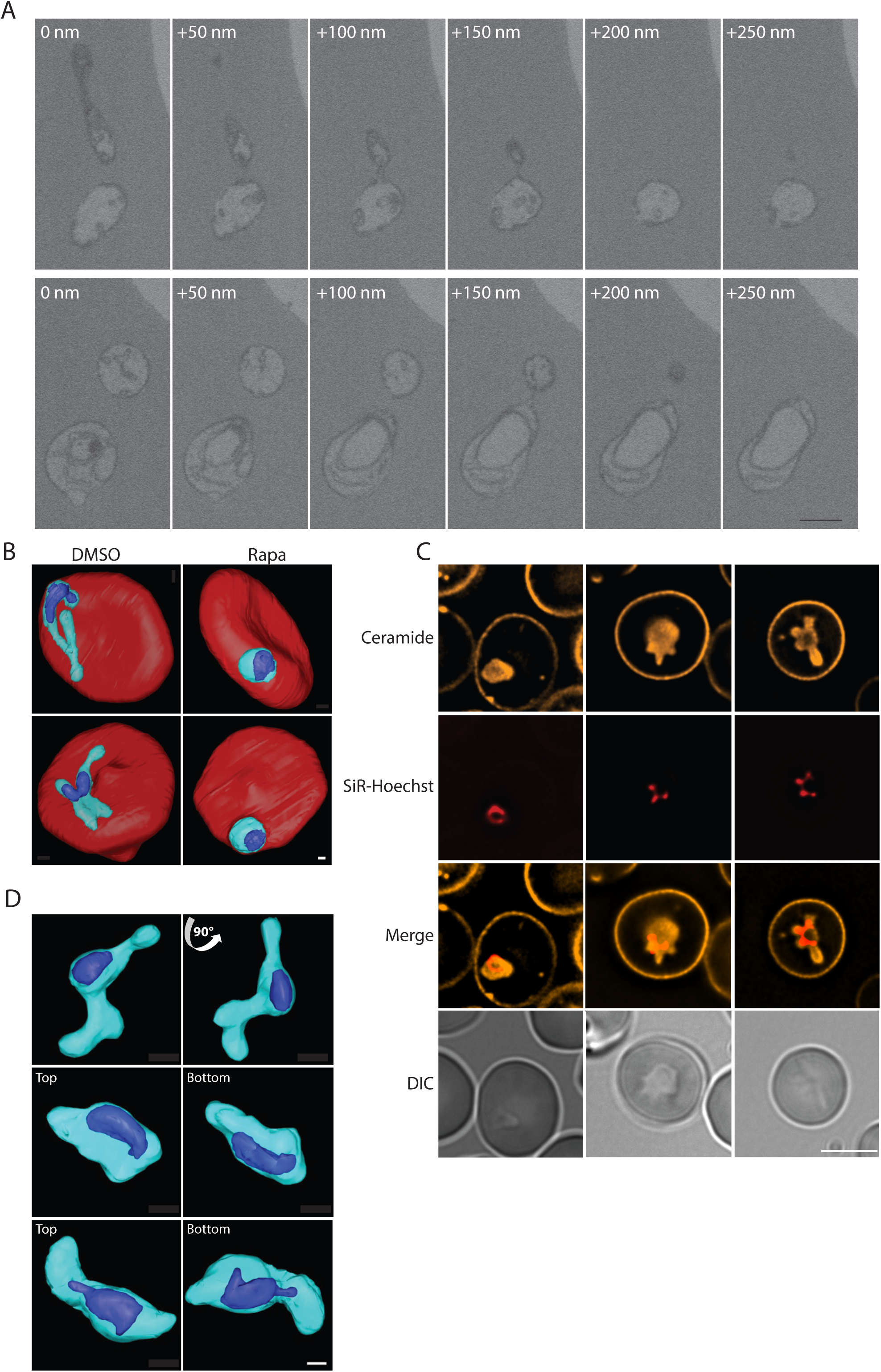
Nuclear morphology in wildtype amoeboid parasites and parasites lacking PV6. **A.** Consecutive SBF-SEM sections, showing the thin connections between limbs of two different parasites. Numbers in the upper left-hand corner of the panels indicate distance between the sections and the scale bar represents 500 nm. **B.** Three-dimensional models from SBF-SEM data of erythrocytes infected with DMSO-treated and rapamycin-treated parasites 2 hours after removal of egress inhibitor. Erythrocytes (red), parasite (cyan) and the nucleus (blue) are highlighted. The scale bar represents 500 nm. **C.** Live-cell fluorescence imaging of infected erythrocytes labeled with C5-Bodipy-ceramide (orange) and SiR-Hoechst (red) 2 hours after removal of egress inhibitor. Note that in the merged imaged, the parasite and the SiR-Hoechst do not overlap perfectly owing to the movement of the parasite during the acquisition of the images. The scale bar represents 5 µm. **D.** Three-dimensional model showing the nucleus (blue) in wildtype parasites (magenta) 20 mins after removal of egress inhibitor. Scale bar represents 500 nm.

**Supplementary Figure 6:**
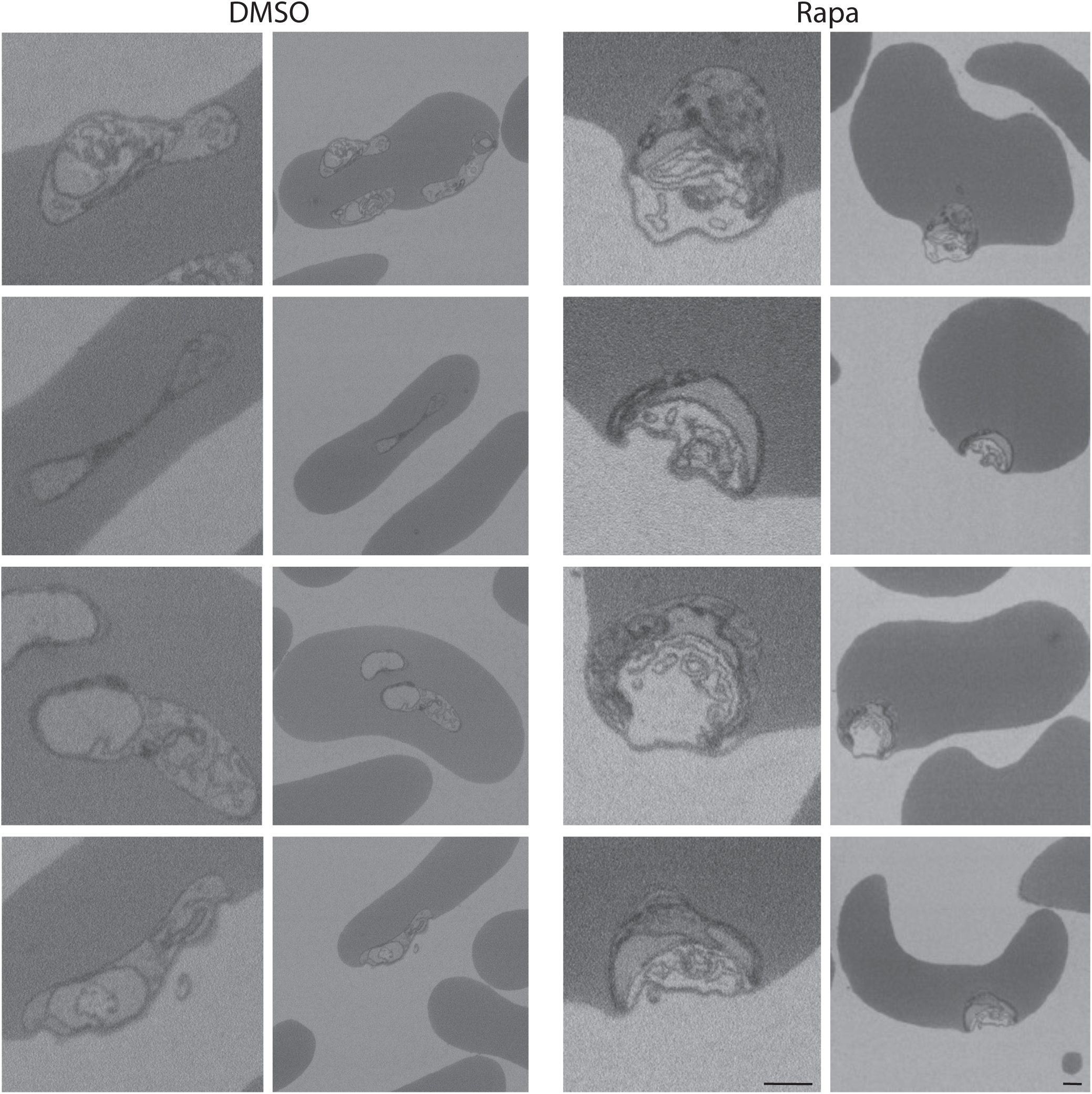
Additional SBF-SEM images of wildtype parasites and parasites lacking PV6. Individual SBF-SEM sections of different erythrocytes infected with wildtype PfBLD529 parasites (left; DMSO) and PfBLD529 parasites lacking PV6 (right; Rapa). In each case, the left-hand panel shows a close view of the parasite and the right-hand panel shows the entire infected erythrocyte. Note the accumulation of membranous whorls next to the parasites lacking PV6. The close views of the parasites in the top row are also shown in Fig 5. Scale bars represent 500 nm.

**Supplementary Figure 7:**
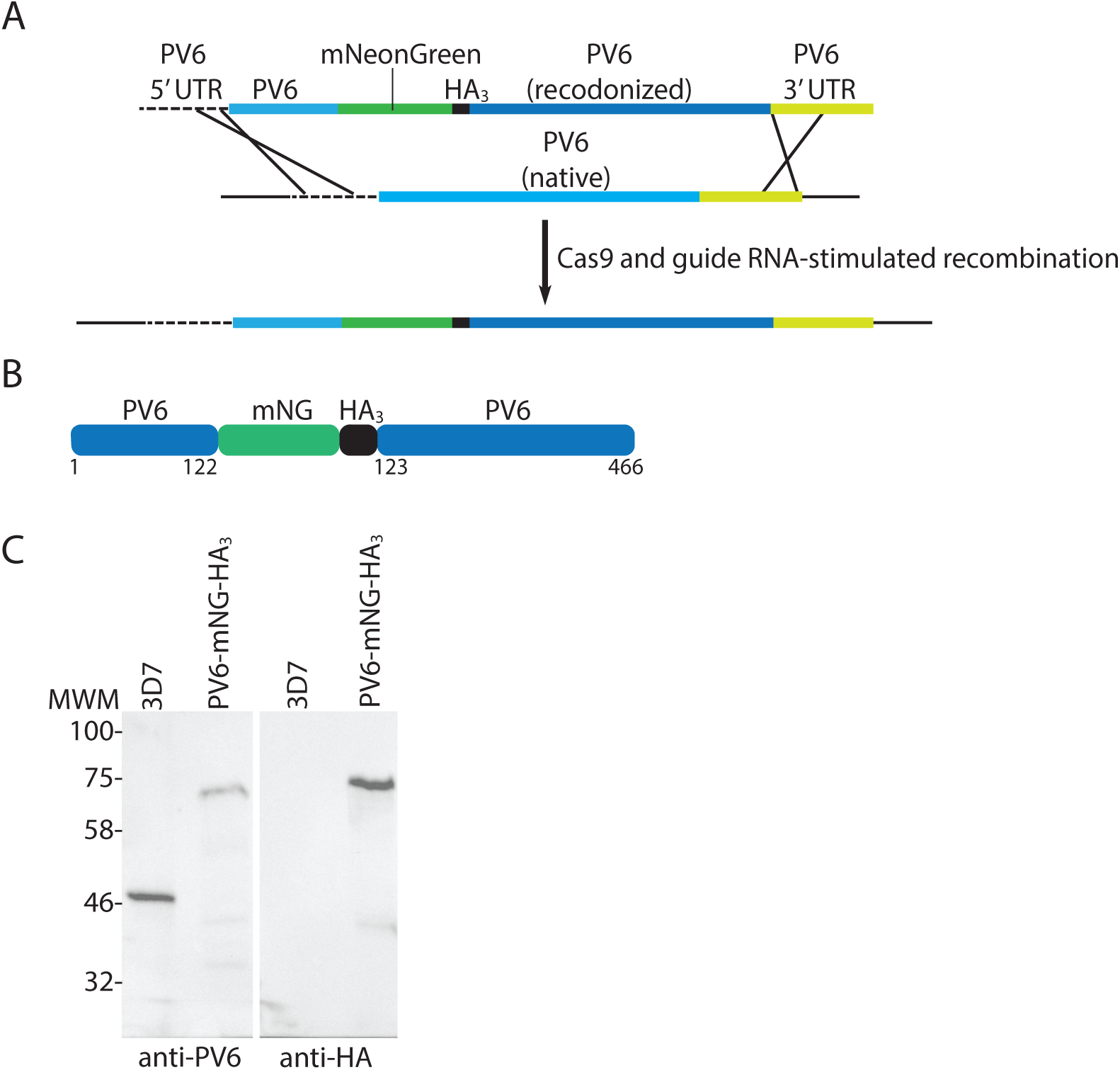
Production of parasites producing a PV6-mNeonGreen fusion. **A.** Outline of the strategy used to produce *Plasmodium falciparum* parasites expressing a mNeonGreen-HA_3_ (mNG-HA_3_)-tagged version of PV6. This strategy is identical to the strategy used by Fréville *et al.*^17^ to modify the *pv6* locus. Cas9 and guide RNA-mediated cleavage of the native PV6 coding sequence promotes its replacement with the fusion gene. The coding sequences of mNeonGreen and the HA_3_-tag are inserted between codon 122 and 123 in the *pv6* gene. **B.** Cartoon showing the expected protein product after insertion of mNG-HA_3_ in PV6 between the AA position 122 and 123. The numbering indicates the amino acid positions in the native PV protein. **C.** Immunoblots on extracts from wild-type (3D7) and transgenic (PV6-mNG-HA_3_) parasites using anti-PV6 (left) and anti-HA (right) antibodies. Expected molecular weights: native PV6 – 46.4 kDa (3D7), PV6-mNG – 72.2 kDa (PV6-mNG). Note the absence of a band representing native PV6 in the extract of PV6-mNG parasites, indicating a very high efficiency of integration.

**Supplementary Figure 8:**
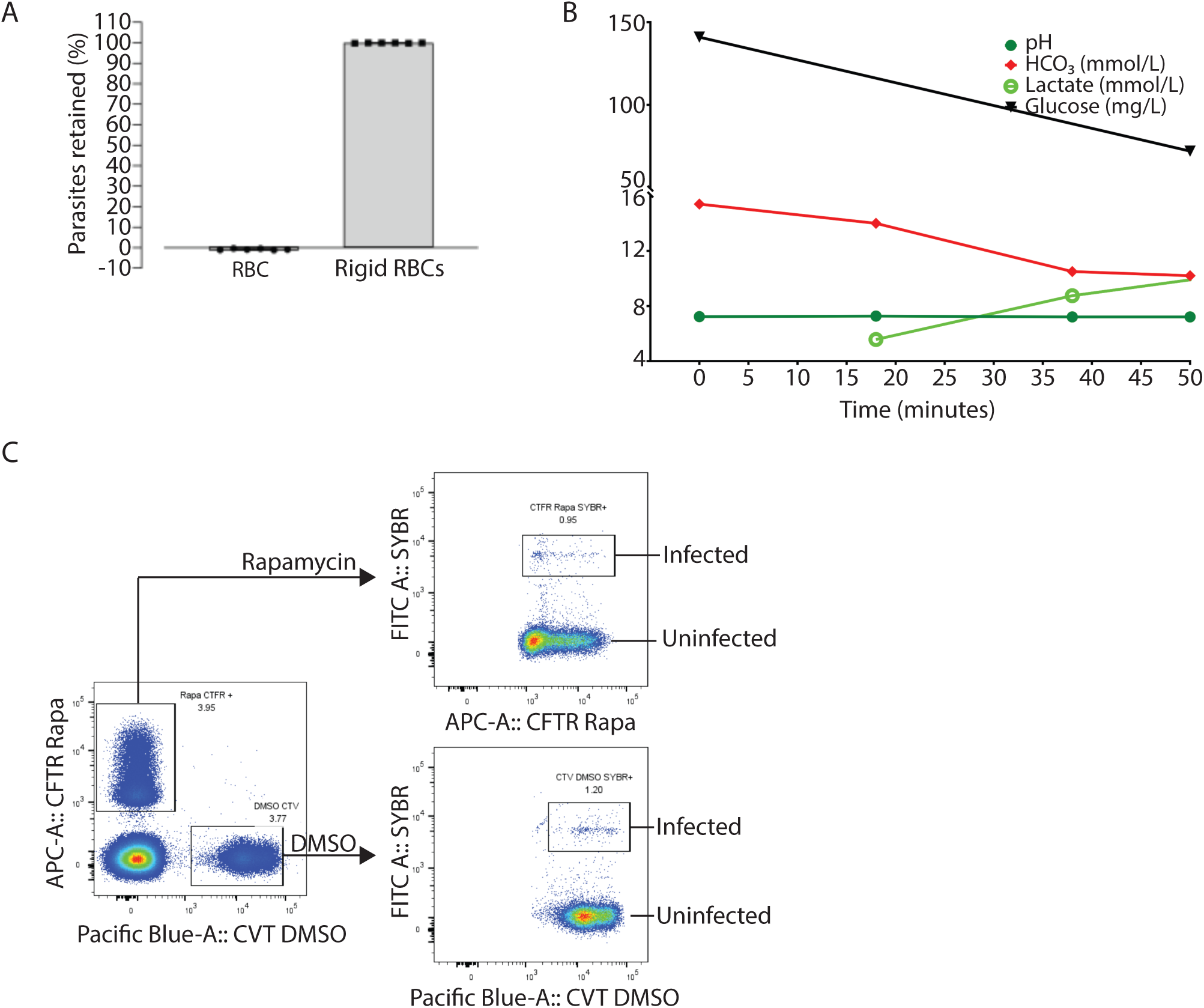
Analysis of samples used for microsphiltration and *ex vivo* spleen perfusion. **A.** Retention of healthy erythocytes (RBCs) and fixed erythrocytes (rigid RBCs) on the columns used for the microsphiltation analysis of erythrocyte infected with wildtype parasites or parasites lacking PV6. **B**. Physiological state of *ex vivo* spleen during the course of the perfusion with infected erythrocytes. The pH and levels of bicarbonate, lactate and glucose were measured periodically to ensure the proper functioning and viability of the spleen. **C.** Gating strategy used for cytometric analysis of blood passaged through *ex vivo* spleen.

